# SARS-CoV-2 ORF7a Drives Mitochondrial Dysfunction via PDK4 Activation and Complex I Inhibition

**DOI:** 10.1101/2025.09.25.678549

**Authors:** Raúl Fernández-Rodríguez, Carmen M. Soto-Jiménez, Rebeca Acín-Pérez, Ana de Lucas-Rius, Luz Marina Sánchez-Mendoza, Blanca D. López-Ayllón, José M. Villalba, José A. Enríquez, María Montoya, Juan J. Garrido, T García-García

## Abstract

SARS-CoV-2 reprograms host metabolism to promote viral replication and evade immune responses. While infection is known to impair mitochondrial function and enhance glycolysis, the role of viral accessory proteins in these alterations remains unclear. Here, we investigate the metabolic impact of the accessory protein ORF7a. Lentiviral expression of ORF7a in human lung epithelial (A549) and monocytic (THP1) cells, coupled with integrated transcriptomic, proteomic, and metabolomic analyses, revealed profound dysregulation of glucose and lipid metabolism. ORF7a impaired oxidative phosphorylation, reducing basal and maximal respiration, inducing mitochondrial depolarization, and increasing reactive oxygen species. Mechanistically, ORF7a upregulated pyruvate dehydrogenase kinase 4 (PDK4), promoting pyruvate dehydrogenase (PDH) complex phosphorylation and suppressing pyruvate oxidation. However, pharmacological PDK4 inhibition did not restore respiration. High-resolution respirometry in frozen samples revealed impaired complex I function, while Blue Native-PAGE demonstrated defective respiratory supercomplex assembly. By linking enzymatic inhibition with structural destabilization, our study uncovers a functional vulnerability of the mitochondrial respiratory chain to viral manipulation. These findings establish ORF7a as a key modulator of host metabolic reprogramming and highlight mitochondrial pathways as potential therapeutic targets in COVID-19.

## Introduction

Severe acute respiratory syndrome coronavirus 2 (SARS-CoV-2), the causative agent of COVID-19, has exemplified profound global impact of zoonotic pathogens. Since its emergence in late 2019, SARS-CoV-2 has led to over 778 million confirmed cases and more than 7 million deaths worldwide (World Health Organization). Although initially characterized primarily as a respiratory illness, accumulating evidence supports that COVID-19 is a complex, multi-system disorder with significant metabolic implications^1,2^. SARS-CoV-2 disrupts systemic and cellular homeostasis, affecting energy metabolism, redox balance, and immune function, thereby contributing to acute respiratory distress syndrome (ARDS), multi-organ failure, and long-term sequelae such as Long COVID^3,4^.

Metabolic reprogramming, the ability of pathogens to hijack host metabolic pathways to support replication, persistence and immune evasion, is a hallmark of many viral infections^5–7^. SARS-CoV-2 profoundly remodels host metabolism by promoting glycolysis, fatty acid biosynthesis, glutaminolysis, and suppressing mitochondrial respiration to create a pro-viral cellular environment^4,8–10^. This metabolic shift is especially pronounced in immune cells, where increased glycolytic flux and impaired oxidative phosphorylation (OXPHOS) impair T-cell function and exacerbate lung epithelial cell death^11^, a phenomenon that may underlie the heightened vulnerability of individuals with metabolic comorbidities such as type 2 diabetes^12^.

A central feature of SARS-CoV-2-induced metabolic remodeling is the shift from mitochondrial respiration to aerobic glycolysis, reminiscent to the Warburg effect observed in cancer cells^11^. This shift is partly driven by the stabilization of hypoxia-inducible factor-1α (HIF-1α), which induces expression of glucose transporters (GLUT1–4), glycolytic enzymes (HK2, PFK1, PKM2, LDHA), and proinflammatory mediators such as interleuckin-1β (IL-1β)^13–15^. Mitochondrial reactive oxygen species (mROS) further stabilizes HIF-1α, creating a feed-forward loop that amplifies inflammation and viral replication. Simultaneously, lactate accumulation suppresses MAVS-dependent antiviral signaling, weakening host immune resposes^16^.

Mitochondria play a central role in cellular energy production, innate immunity and apoptosis^17^. SARS-CoV-2 infection induces profound mitochondrial dysfunction, including cristae disorganization, matrix condensation, membrane depolarization and altered calcium homeostasis^10,18–20^. Electron transport chain (ETC) function is compromised, particularly at complexes I and V, leading to diminished ATP generation, oxidative stress and immune dysregulation^21,22^.

SARS-CoV-2 encodes four structural proteins, sixteen non-structural proteins, and eleven accessory proteins (APs), several of which have been implicated in host metabolic remodeling^23^. Among them, ORF3a is one of the most studied, driving mitochondrial fragmentation, ROS generation, HIF-1α stabilization, and glycolytic activation^14,22,24,25^. ORF9b, ORF9c, and ORF10 also impair mitochondrial respiration, without activating compensatory glycolysis, leading to severe bioenergetic collapse^24^. These findings underscore the ability of APs to subvert host energy metabolism to support viral replication and immune evasion.

In contrast to these APs, ORF7a, a type I transmembrane AP previously implicated in immune evasion and apoptosis, remains poorly characterized in the context of host metabolism. While known to antagonize tetherin (BST2) and downregulate MHC-I expression^26–29^, only limited evidence suggest potential metabolic functions, such as modulation of specific metabolic enzymes^30^. Whether ORF7a directly impairs mitochondrial bioenergetics or contributes to the metabolic dysregulation observed in COVID-19 remains unknown. To our knowledge, this is the first study to systematically investigate ORF7a’s capacity to orchestrate metabolic reprogramming through simultaneous targeting of key mitochondrial pathways.

In this study, we hypothesize that ORF7a modulates host cell metabolism by directly interfering with mitochondrial function. To address this, we ectopically expressed ORF7a in A549 human lung epithelial cells, which serve as a relevant model for respiratory viral infection, and THP1 monocytic cells, representative of innate immune responses, and performed multi-omic profiling alongside functional mitochondrial analyses. We focused particularly on the pyruvate dehydrogenase complex (PDHC), critical metabolic node regulated by pyruvate dehydrogenase kinase 4 (PDK4), whose dysregulation is associated with impaired metabolic flexibility during infection. We also examined the effect of PDK4 inhibition using dichloroacetate (DCA) and assessed the impact of ORF7a on ETC complex I activity.

Our findings reveal a previously unrecognized role for ORF7a in promoting mitochondrial dysfunction through a dual mechanism involving PDK4-mediated PDHC inhibition and direct suppression of complex I. This study positions ORF7a as a critical modulator of host metabolism and highlights mitochondrial pathways as promising therapeutic targets in COVID-19.

## Results

### ORF7a induces metabolic changes affecting glucose and lipid metabolism, nucleotide biosynthesis and redox homeostasis

To investigate the metabolic consequences of ORF7a expression, we performed quantitative proteomic analysis by LC-MS/MS on A549 transduced with ORF7a or control vector. Protein identification using MaxQuant and the Uniprot Homo sapiens database yielded 4.231 proteins with high confidence (**Table S1**). Principle component analysis and hierarchical clustering revealed clear segregation between A549 and A549-ORF7a proteomes (**Figure S1**), underscoring the reproducibility and robustness of the experimental system.

Differential expression analysis identified 244 proteins significantly modulated by ORF7a expression (*q*value < 0.05) in A549-ORF7a compared to control cells (**Figure 1a, Table S2**). Among these, 179 proteins were upregulated, and 65 were downregulated in response to ORF7a expression (**Figure 1a**). Additionally, 80 proteins were uniquely detected in A549-ORF7a cells and 53 exclusively in controls (**Figure 1b, Table S2**) suggesting context-specific protein induction or repression. Gene Ontology (GO) analysis indicated widespread changes in subcellular compartments, including the mitochondria, which accounted for 14% of differentially expressed proteins (**Figure S1c**). Notably, carbamoyl phosphate synthetase 1 (CPS1; log2FC 7.77), a key mitochondrial enzyme involved in nitrogen metabolism, was the most upregulated protein (**Figure 1a**), implicating ORF7a in mitochondrial remodeling and urea cycle perturbation.

**Figure 1.**
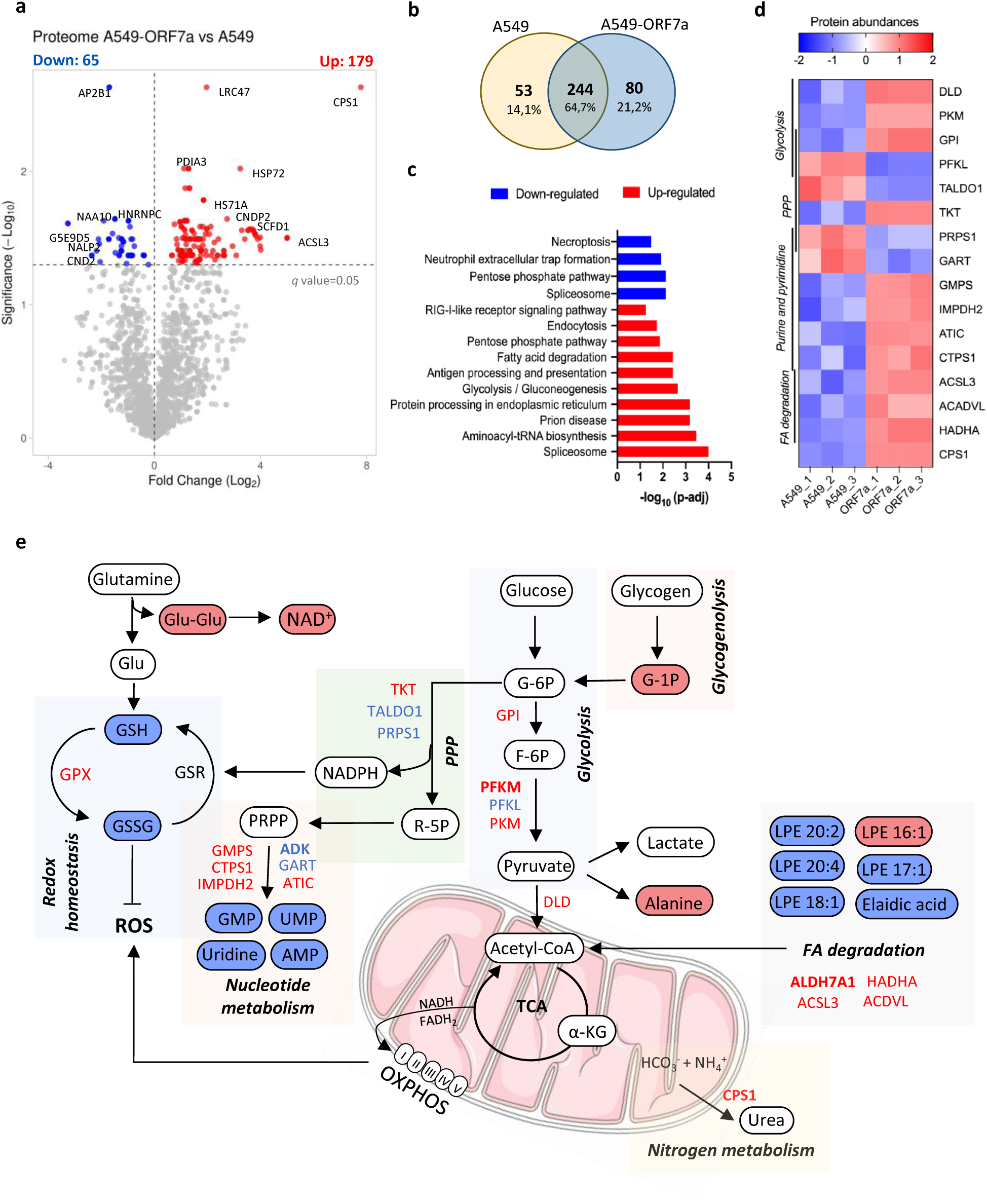
ORF7a-induced metabolic alterations in epithelial cells. a. Volcano plots of protein abundance comparisons between A549-ORF7a and A549 cells. Significant up-regulated proteins were colored in red and down-regulated in blue. b. Overlap of significant differentially expressed proteins identified in A549-ORF7a and control cells A549 c. Enrichment KEGG analysis of up-regulated and down-regulated proteins. d. Heatmaps of changes in protein abundances of components of metabolic pathways in A549 cells expressing ORF7a. e. Schematic representation of key metabolic pathways and metabolites altered upon ORF7a expression in A549 cells. Red-colored metabolites are increased, blue-colored metabolites are decreased, and uncolored intermediates are unchanged or not significantly altered. ORF7a expression induces metabolic reprogramming affecting glycolysis, the pentose phosphate pathway (PPP), the tricarboxylic acid (TCA) cycle, redox homeostasis, and lipid metabolism. Metabolites were identified and quantified via untargeted metabolomics. Red and blue letters represent high and low protein abundance, respectively. Bold red letter represents the present of this protein only in A549-ORF7a and bold blue letter only in A549 cells.

KEGG pathway enrichment analysis revealed that ORF7a modulated a diverse set of pathways including mRNA splicing, protein processing, antigen presentation, and multiple metabolism routes such as glycolysis, fatty acid degradation and the pentose phosphate pathway (PPP) (**Figure 1c**).

We next examined changes in central metabolic enzymes. Several key glycolytic proteins, including glucose phosphate isomerase (GPI), pyruvate kinase (PKM), phosphofructokinase (PFKM) and dihydrolipoamide dehydrogenase (DLD) were significantly upregulated in A549-ORF7a (**Figure 1d-e**), consistent with a glycolytic shift. Enzymes related to fatty acid β-oxidation, such as acyl-CoA dehydrogenase (ACDVL), hydroxyacyl-CoA dehydrogenase trifunctional subunit alpha (HADHA) and acyl-CoA synthetase long-chain family member 3 (ACSL3) were also elevated, suggesting enhanced lipid turnover. Within the PPP, the dowregulation of transaldolase 1 (TALDO1) and phosphoribosyl pyrophosphatase synthase 1 (PRPS1), key for nucleic acid synthesis and purine synthesis, was observed, while transkelotase (TKT) was upregulated, indicating complex modulation of ribose phosphate metabolism **(Figure 1d-e**).

To complement the proteomic data, we conducted untargeted metabolomics profiling using LC-MS in both positive and negative ionization modes to capture hydrophilic and hydrophobic species. Partial least squares-discriminant analysis (PLS-DA) showed clear separation between groups, confirming consistency across replicates (**Figure S2a**). A total of 218 metabolites were significantly altered (IQR <0.40; fold change >1 or <1, p-value < 0.05), of which 29 were annotated (**Table 1**). Volcano plots showed that 107 upregulated and 111 downregulated metabolites in ORF7a-expressing cells (**Figure S2b**).

**Table 1.**
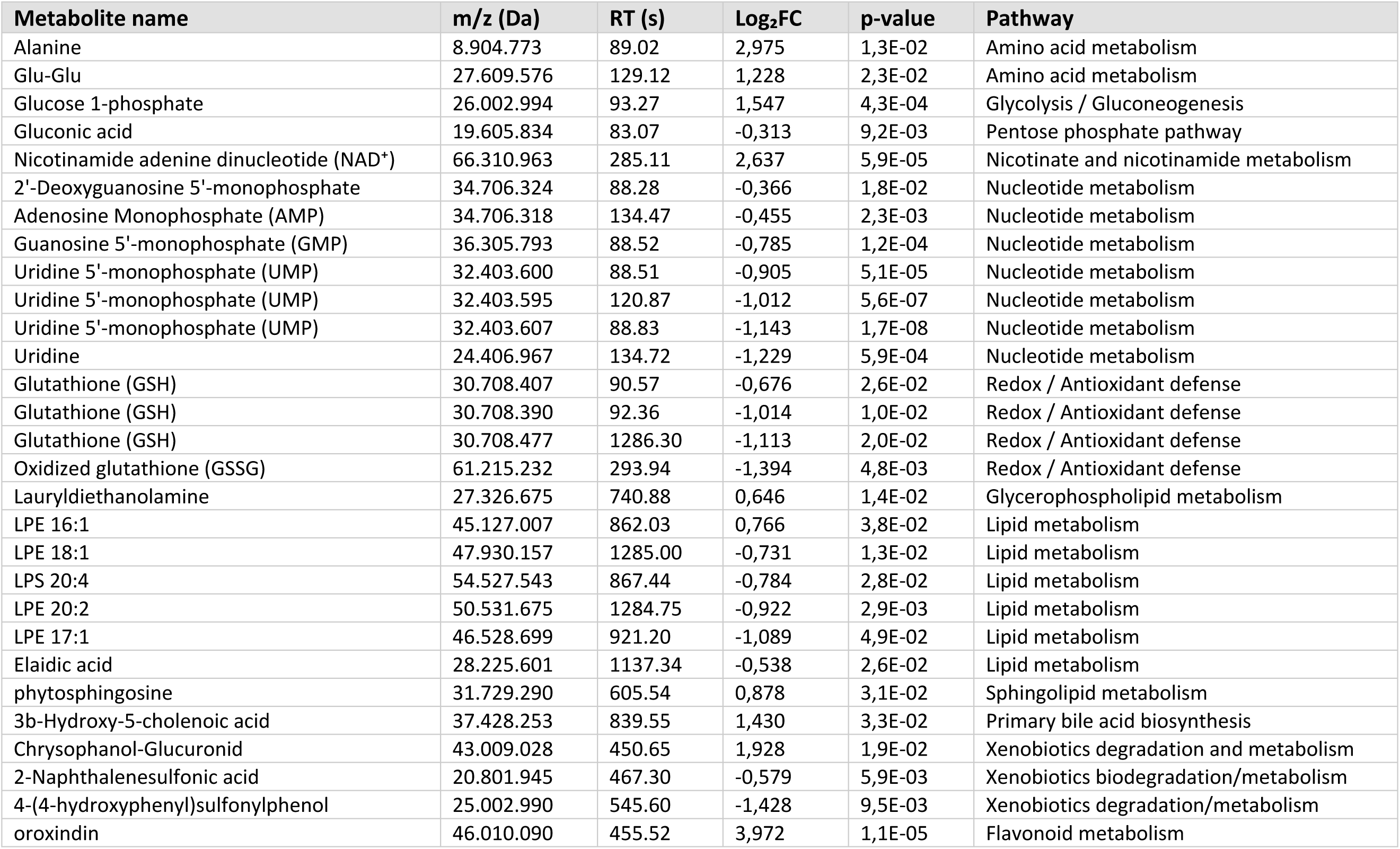
Metabolites significantly altered in A549 cells expressing ORF7a compared with control cells, identified by LC–MS/MS.

Integration of proteomic and metabolomic data revealed significant changes in metabolites involved in glycolysis, nucleotide metabolism, lipid intermediates, amino acid metabolism, and redox homeostasis (**Figure 1e**), glucose 1-phosphate, a key intermediate in glycogenolysis and glycolysis, was significantly elevated (Log_2_FC = 1.54), aligning with increased abundance of glycolytic enzymes. Lipid intermediates, including lysophosphatidylethanolamines (LPE 16:1, 17:1, 20:4), and fatty acids such as elaidic acid were also altered, potentially reflecting membrane remodeling or pro-inflammatory lipid signaling. In contrast, multiple nucleotides monophosphates, including AMP, GMP, UMP, and dGMP were reduced, in agreement with PPP enzyme downregulation (TALDO1, PRPS1). Amino acid metabolism was also affected, as evidenced by elevated Alanine (Log_2_FC = 2.975) suggesting shift in nitrogen and carbon flux. Crucially, several redox-related metabolites, including reduced (GSH) and oxidized glutathione (GSSG), as well as the dipeptide Glu-Glu, were decreased indicating perturbed redox balance and impaired antioxidant capacity (**Figure 1e**).

Together, these results demonstrated that ORF7a expression induces profound metabolic reprogramming in epithelial cells, altering glucose and lipid metabolism, nucleotide biosynthesis and redox homeostasis. This perturbation may contribute to the immunometabolism dysfunction observed in COVID-19 and highlight ORF7a as key driver of host cell metabolic derangement.

### ORF7a enhances glycolysis and promotes HIF-1α stabilization

To functionally validate the metabolic rewiring induced by ORF7a, we performed a Seahorse Glycolysis Stress Test to measure extracellular acidification rate (ECAR), a surrogate measure of glycolytic flux. In A549 lung epithelial cells expressing ORF7a, we observed a significant increase in basal glycolysis compared to control cells (**Figure 2a–b**). To assess whether this effect extend to immune cells, TPH1 monocytes were transduced with ORF7a and differentiated to macrophage-like cells (MΦ). Notably, glycolytic activation was more pronounced in THP1-MΦ expressing ORF7a, which shower enhanced ECAR following glucose and oligomycin injection, indicating more robust glycolytic flux in these immune cells **(Figure 2c-d**).

**Figure 2.**
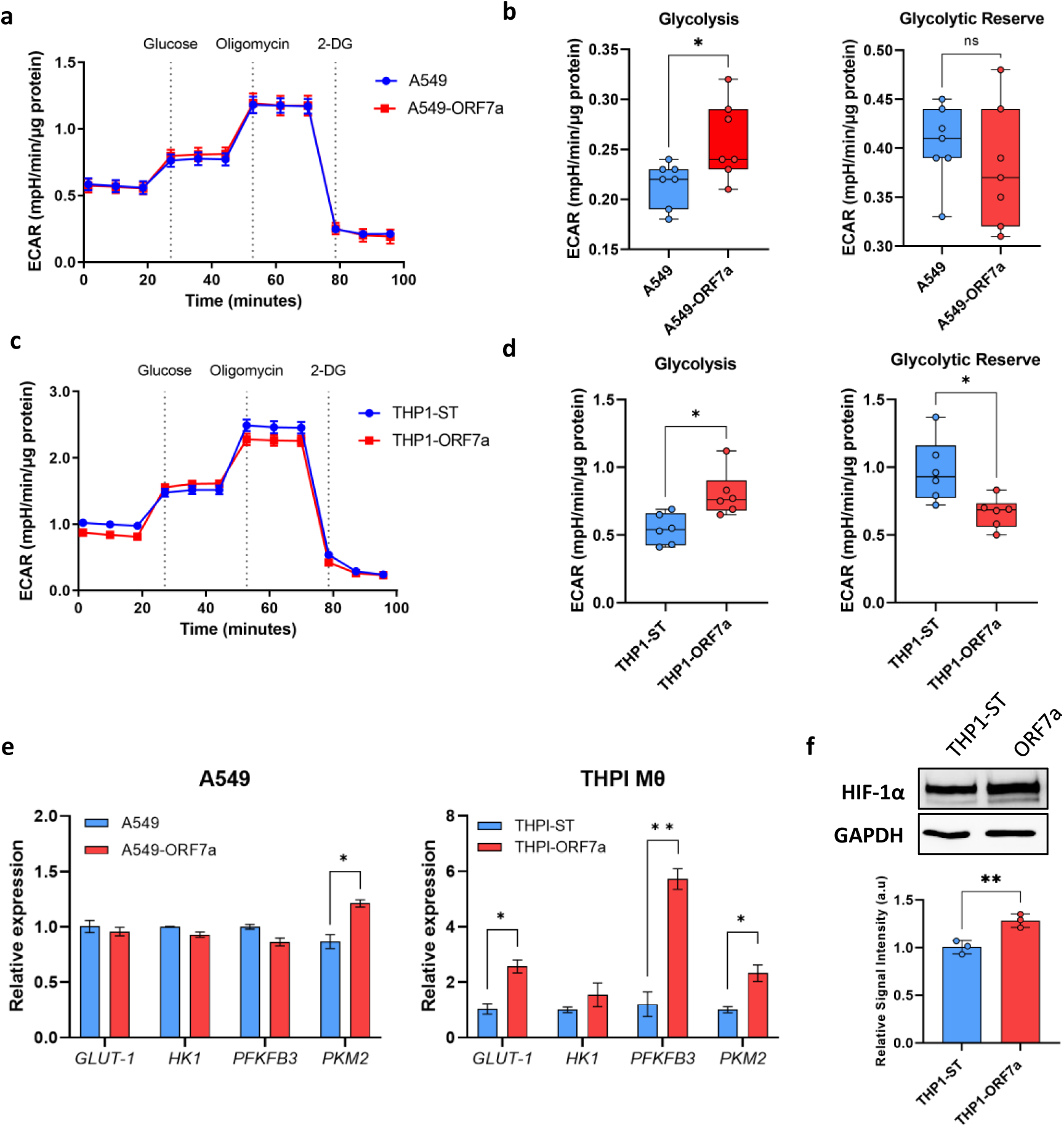
ORF7a enhances glycolytic activity in epithelial and macrophage-like cells. **a-d.** Seahorse glycolysis stress test profile measuring extracellular acidification rate (ECAR) in glucose-free media followed by sequential addition of glucose, oligomycin and 2-deoxy-D-glucose (2-DG) used to determine glycolytic metrics in A549 transduced cells (a-b) or THP1 transduced cells (c-d) using Seahorse XFe24 analyzer. Parameters of Glycolysis and Glycolytic reserve calculated by the Wave program in A549 transduced cells (b) and THP1 transduced cells (d). **e**. Relative mRNA expression of main glycolytic genes in A549 and THP1 derived macrophage-like cells (THP1 MƟ) expressing ORF7a measured by qPCR. **f**. Western blot showing HIF-1α stabilization in THP1-ORF7a cells were treated with CoCl2 for 6 h. Error bars represent the mean ± SEM. Statistical significance is given as follows: *p < 0.05, **p < 0.01 and ***p < 0.00.

To investigate the transcriptional basis of this glycolytic phenotype, we quantified the expression of key metabolic regulators. *GLUT-1, PFKFB3* and *PKM2*, are known to be upregulated during SARS-CoV-2 infection, contributing to the metabolic reprogramming observed in COVID-19 patients^11^. Consistent with our proteomic data, we found that *PKM2* expression, was significantly increased in both A549 and THP1 cells expressing ORF7a (**Figure 2e**). Additionally, *GLUT-1* and *PFKFB3* were selectively upregulated in THP1-ORF7a cells, correlating with the stronger glycolytic response in this lineage (**Figure 2e**). These enzymes regulate rate-limiting steps in glucose uptake and flux through glycolysis, supporting the notion that ORF7a enhances glycolytic metabolism, particularly in innate immune cells.

Given the coordinated upregulation of glycolytic genes, we hypothesized that ORF7a may engage HIF-1α, a central transcriptional driver of glycolysis under both hypoxia and inflammatory conditions. To test this, we assessed HIF-1α protein stabilization following treatment with CoCl₂, a hypoxia-mimetic agent. While CoCl₂ induced HIF-1α in both control and ORF7a-expressing THP1-MΦ cells, the response was markedly amplified in the presence of ORF7a (**Figure 2f**), suggesting that ORF7a enhances glycolysis, at least in part, through HIF-1α-dependent transcriptional programs.

These finding are consistent with previous reports of HIF-1α activation in SARS-CoV-2-infected cells and tissues, where it promotes both metabolic reprogramming and inflammatory signaling^11,21^. Furthermore, APs such as ORF3a have been shown to trigger HIF-1α stabilization^14^. Our data suggest that ORF7a may function via a similar mechanism, acting through HIF-1α to amplify glycolytic flux and support a pro-viral cellular environment.

### ORF7a impairs mitochondrial respiration and induces oxidative stress in lung epithelial cells

Given the observed glycolytic shift induced by ORF7a, we next investigated whether mitochondrial oxidative phosphorylation (OXPHOS) was functionally compromised. Using the Seahorse MitoStress Test, we measured oxygen consumption rate (OCR), a key indicator of mitochondrial respiration. ORF7a expression led to a significant reduction in basal respiration and ATP-linked respiration in A549 cells, indicating impaired mitochondrial energy production (**Figure 3a–b**). Additionally, both maximal respiration and spare respiratory capacity were markedly decreased, suggesting that ORF7a limits the cells’s ability to adapt to metabolic stress (**Figure 3b**). To determine whether these defects were associated with structural alterations, we performed transmission electron microscopy. No significant changes in mitochondrial length or cristae density were detected between ORF7a-expressing and control cells (**Figure 3c-d**). However, MitoView 633 staining followed by flow cytometry revealed a significant increase in the proportion of low-activity mitochondria in A549-ORF7a cells (> 10%), consistent with impaired OXPHOS function (**Figure 3e–f**). Importantly, these functional defects were not attributable to loss of mitochondrial content of downregulation of electron transport chain (ETC) components, as total mitochondrial protein levels and ETC subunits remained unchanged (**Figure S3**).

**Figure 3.**
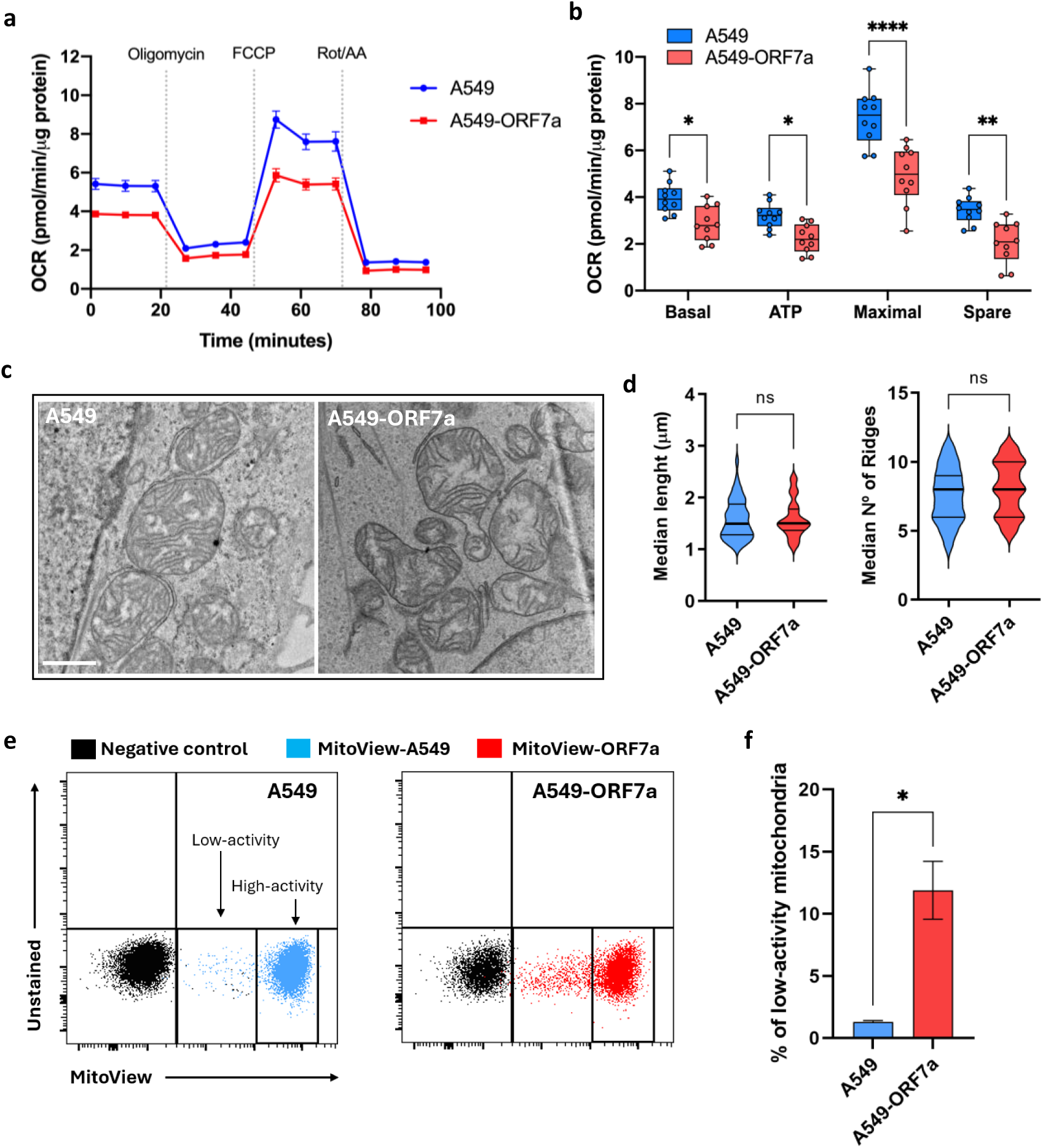
Effects of SARS-CoV-2 ORF7a on mitochondria function. **a.** MitoStress Test performed on Seahorse XFe24 analyzer. Canonical mitochondrial inhibitors injected sequentially as labelled, and the oxygen consumption rate (OCR) was calculated and normalized to quantity of protein. **b**. MitoStress Test parameters calculated from OCR data. **c**. Analysis by transmission electron microscopy (TEM) of mitochondrial appearance in A549-ORF7a cells. Scale bar indicates 1 µm. **d**. Median of mitochondria longest diameter and median number of ridges per mitochondrion evaluated from TEM micrographs of each cell line. In both cases, at least 50 mitochondria were analyzed. e. Data representative of flow cytometry showing high and low-activity mitochondria in A549 and A549-ORF7a cells stained with MitoView 633. **f**. Quantification of the percentage of low-activity mitochondria in A549-ORF7a cells stained with MitoView 633. Data are represented as mean ± SD (n = 4). Statistical significance is given as follows: *p < 0.05, **p < 0.01, ***p < 0.001 and ****p < 0.0001.

Mitochondria are a major source of ROS, and excessive ROS production can lead to oxidative damage and impaired mitochondrial function (**Figure 4a**)^31^. To explore whether the impaired respiratory function was accompanied by increased oxidative stress, a hallmark of mitochondrial dysfunction, we analyzed redox-related metabolites. Untargeted metabolomics revealed a significant reduction in glutathione species (GSH, GSSG) and the dipeptide Glu-Glu (**Figure 4a**) indicating compromised redox buffering capacity. Consistently, intracellular ROS levels measured by H₂DCFDA staining and flow cytometry were significantly elevated in A549-ORF7a cells (**Figure 4b**), suggesting that ORF7a expression promotes oxidative stress. To assess whether the antioxidant defense system was affected, we analyzed the expression of ROS-scavenging enzymes. Transcriptomic profiling revealed altered expression of key antioxidant regulators (**Figure 4c**), including down-regulation of peroxiredoxin 2 (PRDX2), a highly efficient peroxide scavenger, and upregulation of glutathione peroxidase 1 (GPX1). These transcriptomic changes were validated by qPCR, which confirmed increased *GPX1* expression in both A549- and THP1-ORF7a cells (**Figure 4d**), suggesting a compensatory antioxidant response to increased oxidative burden.

**Figure 4.**
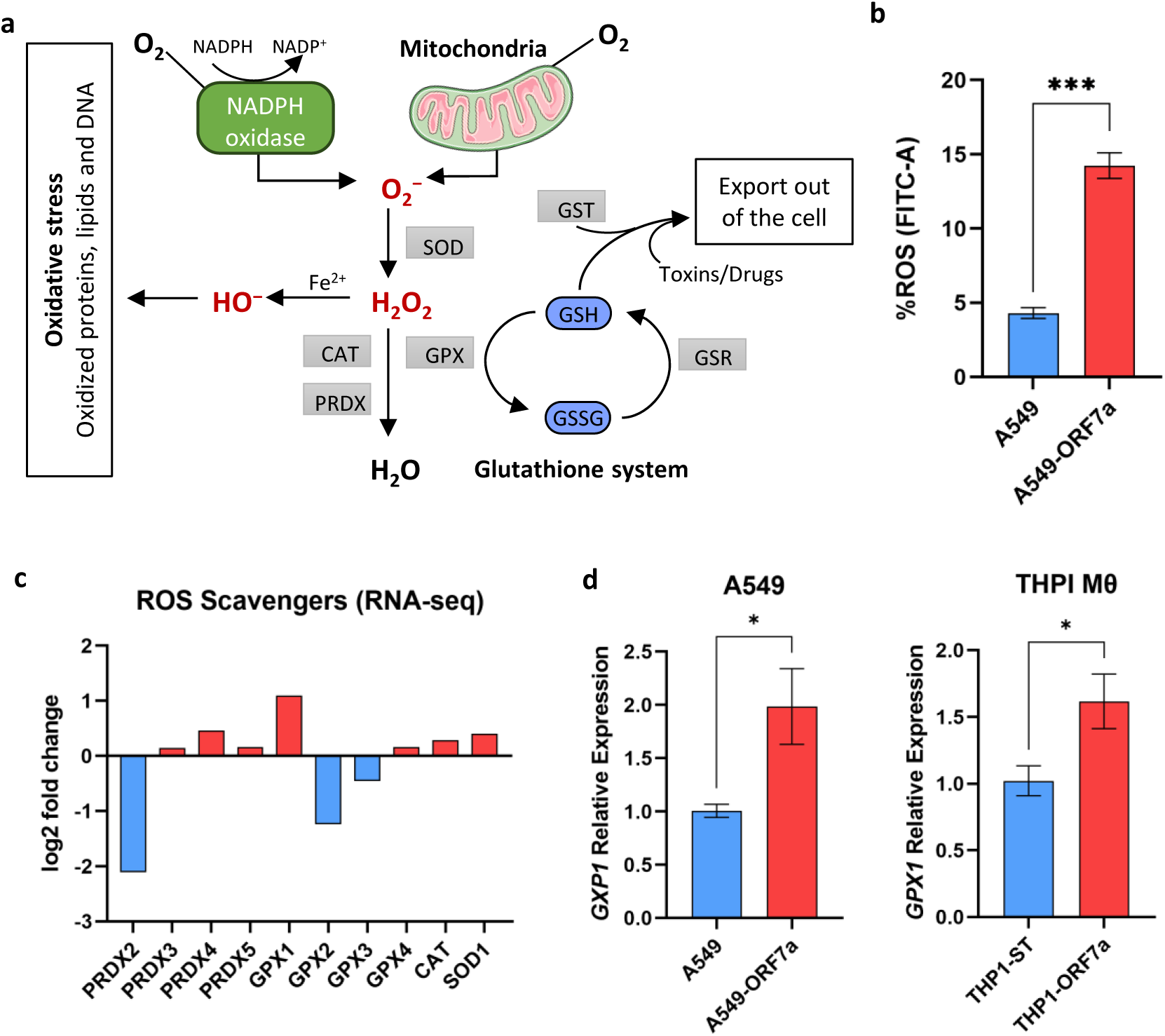
ORF7a impairs ROS homeostasis by altering antioxidant defense systems. **a.** Schematic representation of ROS generation and scavenging. ROS scavengers are antioxidant enzymes that neutralize ROS by directly reacting with and accepting electrons from ROS. When ROS production exceeds the capacity of scavenging systems, ROS accumulate excessively, leading to oxidative stress and damaging cellular components such as proteins, lipids and nucleotides. Cells counteract this imbalance through various antioxidants systems that targeted specific ROS species, helping to prevent pathological ROS levels and to repair oxidative damage. Key antioxidant enzymes include superoxide dismutase (SOD), catalase (CAT), peroxiredoxins (PRX), glutathione peroxidase (GPX) and glutathione reductase (GSR). **b**. ROS production measured by flow cytometry. **c**. Log2 fold changes of significant altered ROS scavengers identified in the RNA-seq. **d**. Relative mRNA expression of *GXP1* in A549 and THP1 derived macrophage-like cells expressing ORF7a, measured by qPCR. Error bars represent the mean ± SEM. Statistical significance is given as follows: *p < 0.05, **p < 0.01 and ***p < 0.001

Taken together, our multi-omics and functional analyses demonstrated that ORF7a expression induces a robust oxidative stress phenotype, likely driven by mitochondrial dysfunction and an inadequate antioxidant response, not through depletion of mitochondrial mass or ETC components, but likely via impairment of mitochondrial function and ROS homeostasis. These findings support a model in which ORF7a contributes to COVID-19 pathophysiology by promoting immunometabolic dysfunction through mitochondrial stress.

### ORF7a Disrupts Mitochondrial Metabolism via PDK4-Mediated Inhibition of the Pyruvate Dehydrogenase Complex

Mitochondria play a central role in coordinating metabolic flux between glycolysis, fatty acid oxidation, and the tricarboxylic acid (TCA) cycle, and has been identified as key targets of SARS-CoV-2 activity. To examine how ORF7a affects mitochondrial protein expression, we cross-referenced differentially expressed proteins (DEPs) from our proteomic dataset with the curated MitoCarta 3.0 database. We identified 19 mitochondrial DEPs in A549 cells expressing ORF7a (**Figure 5a**). To complement this, we compared these findings with RNA-seq data from ORF7a-expressing A549 cells^30^, identifying 20 mitochondrial genes differentially expressed at the transcript level (fold change > 2; p < 0.05). Strikingly, only CPS1 overlapped between the proteomic and transcriptomic datasets, suggesting that many mitochondrial changes may be regulated post-transcriptionally.

**Figure 5.**
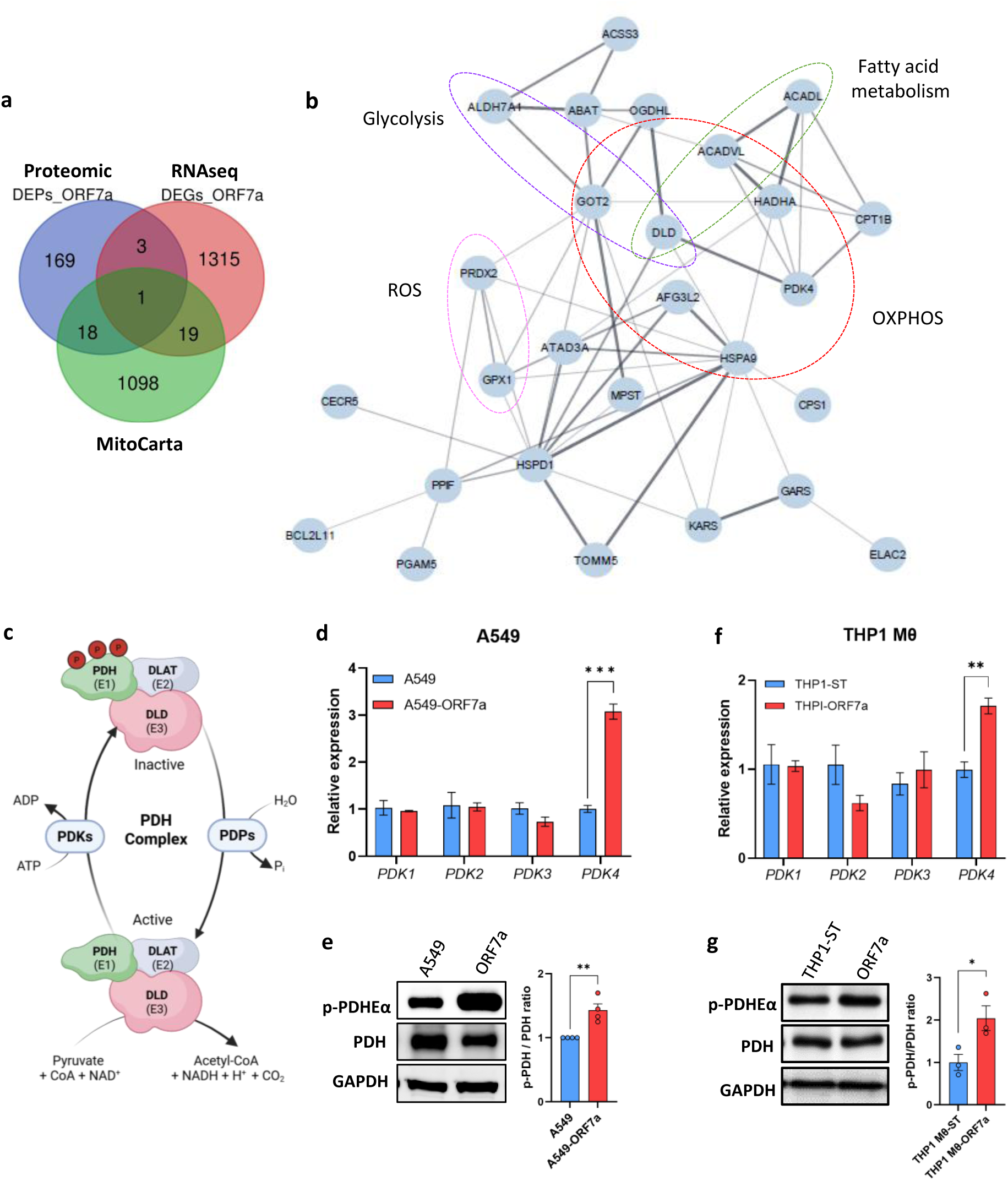
ORF7a increases the expression of PDK4 and phosphorylation of PDHE1α. **a**. Venn diagram comparing all Mitocarta3.0 entries with differential expressed proteins (DEPs) and differential expressed genes (DEGs) from A549 cells expressing ORF7a. **b**. STRING interaction network of mitochondrial DEPs and DEGs highlighting enrichment of Hallmark pathways. **c**. Schematic representation of PDH complex regulation. **d-g**. Effect of ORF7a on mRNA expression of PDK isoforms (PDK1, PDK2, PDK3 and PDK4) and phosphorylation of PDHE1α (Ser300) using a phospho-specific PDHE1α antibody in pulmonary epithelial A549 cells (**d,e**) and THP1 derived macrophage-like cells (THP1 MƟ) (**f,g**). Values are means ± SEM. n = 3 per group. *p < 0.05, **p < 0.01 and ***p < 0.001 vs. control cells.

To identify potential regulatory hubs, we constructed a protein-protein interaction (PPI) network from the union of mitochondrial DEPs and DEGs (**Figure 5b**). This analysis revealed clusters of interacting proteins involved in OXPHOS, glycolysis, fatty acid metabolism and ROS regulation. Among these, DLD emerged as central node. DLD is the E3 subunit of the pyruvate dehydrogenase (PDH) complex (**Figure 5c**), a mitochondrial multi-enzyme complex that catalyzes the oxidative decarboxylation of pyruvate. PDH complex controls the conversion of pyruvate to acetyl-CoA linking glycolysis to the TCA cycle and mitochondrial respiration^32–34^.

To investigate whether PDH function is compromised by ORF7a, we analyzed the expression of pyruvate dehydrogenase kinases (PDKs), which negatively regulate the PDH complex by phosphorylation of its E1α subunit (**Figure 5c**). PDK isoforms are known to be dysregulated in pathological conditions such as diabetes, obesity, and cancer inhibit the activity of the PDH complex^35–37^.

ORF7a expression significantly increased *PDK4* mRNA levels in A549 cells (**Figure 5d**), while expression levels of *PDK1*, *PDK2* and *PDK3* remained unchanged. We next examined the phosphorylation status of PDHE1α at serine 300, well-characterized site of PDH inhibition^33,38^. Western blot analysis revealed robust phosphorylation of PDHE1α in ORF7a-expressing A549 cells compared to controls (**Figure 5e**), indicating functional inactivation of the PDH complex. Similar results were observed in THP1-MƟ (**Figure 5f-g**), in which ORF7a also induced both PDK4 upregulation and increased PDHE1α phosphorylation.

Collectivity, these findings demonstrate that ORF7a induces PDK4 upregulation and consequent inhibition of PDH activity, thereby redirecting pyruvate flux away from mitochondrial oxidation. Interestingly, while PDH inhibition would typically lead to pyruvate accumulation and lactate production, metabolomic analysis revealed increased levels of alanine (**Figure 1e**, **Table 1**), with no significant accumulation of pyruvate itself. This suggests an alternative metabolic route in which pyruvate is preferentially transaminated to alanine-by-alanine aminotransferase, rather than entering the mitochondria. Such a pathway may serve as a compensatory mechanism to avoid redox stress associated with pyruvate buildup and maintain cytosolic NAD⁺/NADH balance under conditions of impaired mitochondrial oxidation

### ORF7a impairs mitochondrial respiration by selectively disrupting complex I activity

To assess whether the mitochondrial dysfunction induced by ORF7a could be reverse, we treated cells with dichloroacetate (DCA), a well-characterized inhibitor of PDK4. By suppressing PDK4, DCA reactivates the PDH complex, enhancing the conversion of pyruvate to acetyl-CoA and promoting its entry into the TCA cycle^39^. In THP1 control cells, DCA treatment led to a marked decrease in phosphorylation of PDHE1α at Ser300 by western blot. DCA treatment markedly reduced PDHE1α phosphorylation levels in, confirming effective PDH reactivation (**Figure 6a**). This was accompanied by robust increase in maximal respiration measured by Seahorse MitoStress Test (**Figure 6b**), consistent with enhances oxidative capacity upon PDH reactivation.

**Figure 6.**
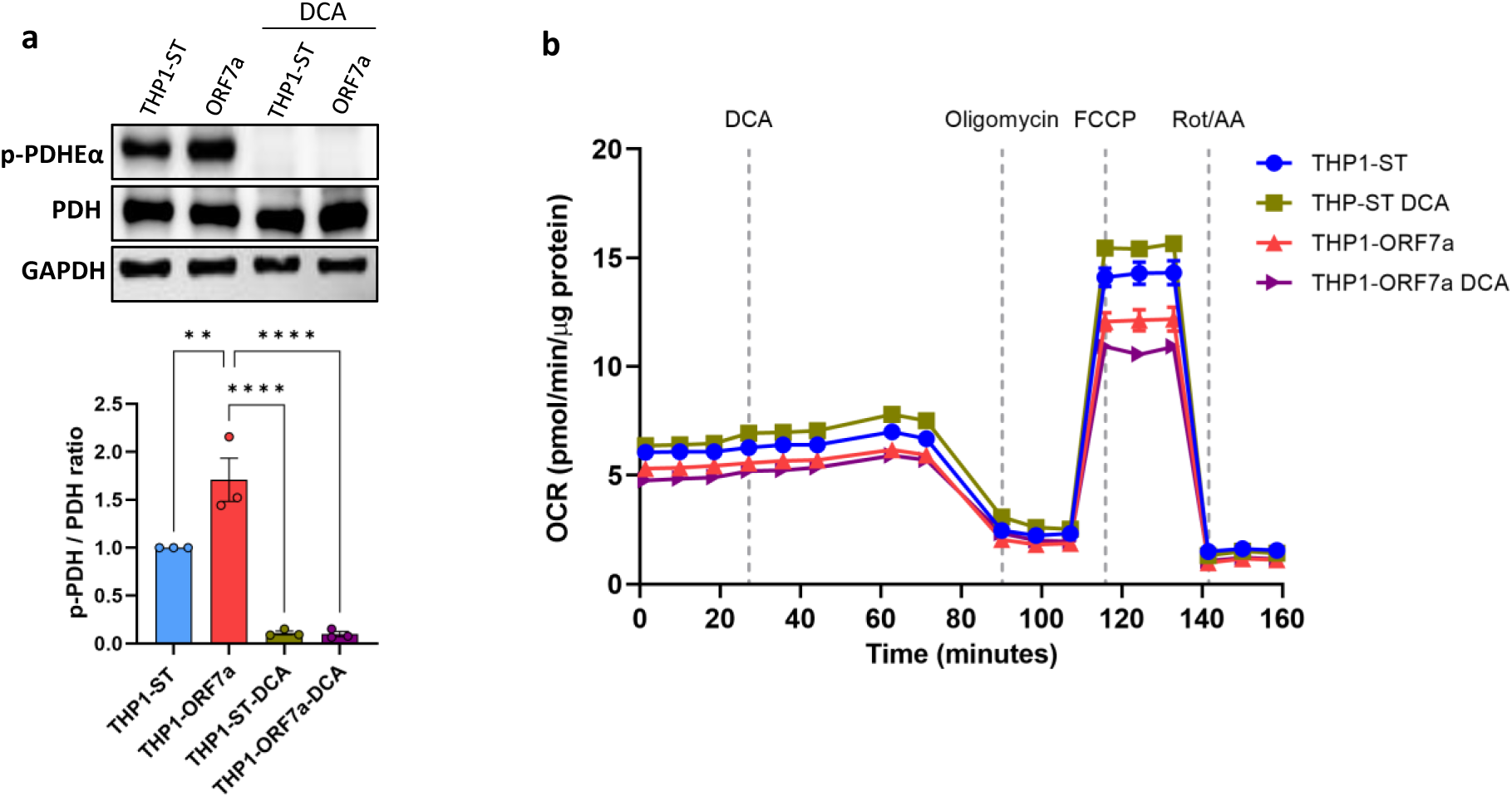
ORF7a impairs mitochondrial respiration and prevents recovery by DCA-mediated PDH reactivation in THP1 cells. **a**. Western blot analysis of phosphorylated PDHE1α (Ser300), total PDH, and GAPDH in control and ORF7a-expressing cells with or without DCA treatment. Quantification of p-PDHE1α/PDH ratio shows a significant decrease upon DCA treatment in both conditions, indicating effective PDH activation. **b**. Mitochondrial respiration measured using the Seahorse MitoStress Test in THP1 and

Surprisingly, in ORF7a-expressing THP1 cells, DCA failed to restore mitochondrial respiration. Instead, we observed a further reduction in maximal OCR after DCA treatment (Figure 6b), indicating that the mitochondrial impairment caused by ORF7a is not solely due to PDH inhibition. These findings suggest the involvement of additional defects affecting mitochondrial bioenergetics downstream or independent of PDH regulation.

To identify which components of the ETC were affected, we performed a series of substrate-driven respiration assays in permeabilize THP1 cells using a Seahorse XF analyzer. In the first assay (**Figure 7a**), we assessed complex I–dependent respiration by adding NADH, followed by antimycin A to inhibit complex III, and then TMPD/ascorbate to evaluate complex IV activity. THP1 cells expressing ORF7a exhibited a significantly reduced OCR in response to NADH, compared to control cells, while complex IV-mediated respiration (TMPD/ascorbate-driven) remained unchanged, suggesting that complex IV activity remained intact.

**Figure 7.**
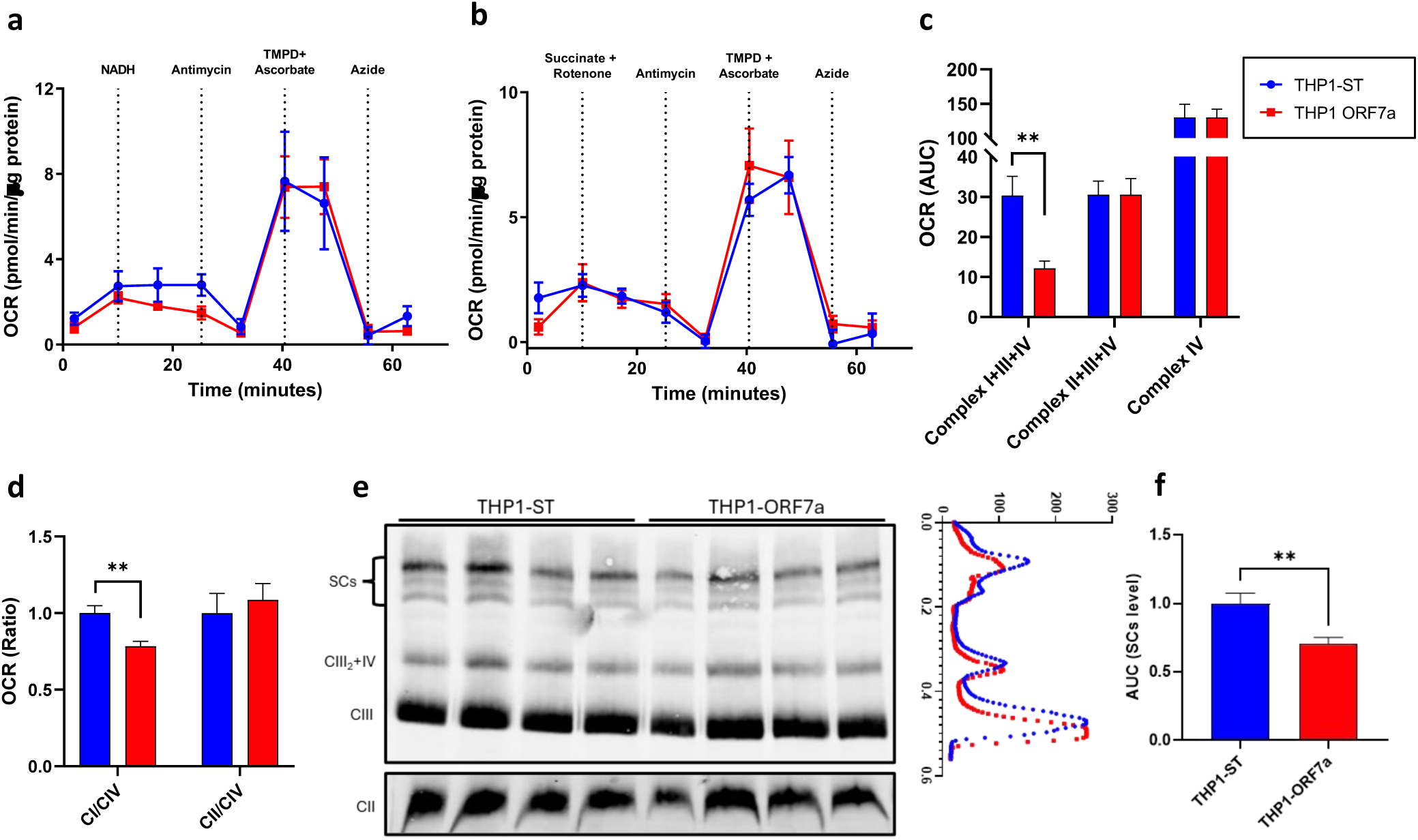
ORF7a expression impairs mitochondrial complex I–dependent respiration. **a**. OCR in permeabilized THP1 cells expressing ORF7a or control vector (THP1-ST), following sequential addition of NADH (complex I substrate), antimycin A (complex III inhibitor), TMPD/ascorbate (complex IV substrate), and sodium azide (complex IV inhibitor) measured by Seahorse XF24 **b**. OCR traces of permeabilized THP1 cells following addition of succinate (complex II substrate) in the presence of rotenone (complex I inhibitor), followed by antimycin A, TMPD/ascorbate, and azide. **c**. Quantification of the area under the curve (AUC) from Seahorse traces. Data are presented as mean ± SEM. p < 0.05 by unpaired t-test (n = 8). **d**. Quantification of the OCR ratio CI/CIV and CII/CIV OCR in THP1 mitochondrial fraction measured by Seahorse XF96. Data are presented as mean ± SEM. p < 0.05 by unpaired t-test (n = 4). **e**. Representative blot (left) and intensity profile (right) showing ETC complexes assembly by BN-PAGE in THP1 mitochondrial fraction. **f**. Quantification of the AUC from Supercomplexes (SCs) level at intensity profile. Data are presented as mean ± SEM. p < 0.05 by unpaired t-test (n ≥ 4).

To evaluate complex II–dependent respiration, we used a second assay (**Figure 7b**) in which succinate was added in the presence of rotenone (to block complex I). Under these conditions, no significant differences in OCR were observed between ORF7a-expressing and control cells, indicating that complex II–mediated respiration was preserved. Quantification of the area under the curve (AUC) from both Seahorse assays (**Figure 7c**) confirmed a selective reduction in complex I-dependent respiration in ORF7a-expressing cells, while complex II and complex IV remained unaffected.

To strengthen these findings, we performed the same substrate-driven respiration assays in isolated mitochondria form control and ORF7a-expressing THP1 cells (**Figure 7d**). The results fully recapitulated those obtained in permeabilized cells: a pronounced impairment of complex I–dependent respiration, while complexes II and IV remained unaffected. Performing the assays in isolated mitochondria rules out the possibility that the observed defect was due to altered NADH availability or to the activity of alternative NADH-consuming enzymes, confirming that ORF7a selectively impairs complex I–mediated electron transfer.

To further investigate whether the respiratory defect induced by ORF7a was associated with alterations in the structural organization of the ETC, we analyzed the assembly of mitochondrial supercomplexes by Blue Native PAGE (BN-PAGE) (**Figure 7e**). Control mitochondria displayed the expected pattern of high–molecular weight supercomplexes (SCs), including the respirasome (I+III_2_+IV_n_). In contrast, mitochondria isolated from ORF7a-expressing THP1 cells exhibited a marked reduction in the abundance of fully assembled supercomplexes, particularly complex I. Densitometric quantification confirmed a significant decrease in the proportion of complex I incorporated into supercomplexes (**Figure 7f**), suggesting that ORF7a impairs not only the enzymatic activity of complex I but also its supramolecular assembly within the respiratory chain.

Taken together, these findings demonstrate that ORF7a selectively impairs complex I activity, independently of PDH inhibition, resulting in defective electron transfer and reduced respiratory capacity. The additional observation that ORF7a disrupts the incorporation of complex I into supercomplexes provides mechanistic insight for the loss of OXPHOS efficiency, establishing complex I dysfunction as a key hallmark of ORF7a-expressing cells.

## Discussion

Our study uncovers a multifaceted role for the SARS-CoV-2 accessory protein ORF7a as a modulator of host cell metabolism and mitochondrial function. Through integrated proteomic, transcriptomic, metabolomic, and functional assays, we demonstrate that ORF7a reprograms central carbon metabolism, suppresses OXPHOS, induces oxidative stress, and selectively impairs complex I of ETC. These findings identify ORF7a as a critical driver of metabolic remodeling in SARS-CoV-2 pathogenesis.

Quantitative proteomics and untargeted metabolomics revealed significant alteration in pathways involving glycolysis, lipid turnover, nucleotide biosynthesis, and redox balance (**Figure 1**). ORF7a expression was associated with upregulation of key glycolytic enzymes (PKM, PFKM, GPI) and accumulation of glucose 1-phosphate, consistent with the enhanced glycolytic flux (**Figure 1d-e**). Seahorse extracellular flux analysis confirmed this phenotype (**Figure 2a-d**), showing elevated ECAR, particularly in macrophage-like THP1 cells, suggesting that ORF7a may preferentially reprogram innate immune cells toward glycolysis. These findings are consistent with previous studies reporting metabolic reprogramming toward glycolysis in SARS-CoV-2 infection ^4,8,11,40–42^. Mechanistically, this glycolytic switch was linked to stabilization of HIF-1α and transcriptional upregulation of GLUT-1, PFKFB3, and PKM2 (**Figure 2e-f**), in line with previous reports of HIF-1α–driven metabolic adaptation during SARS-CoV-2 infection^11,14,21^ and suggest that ORF7a, like ORF3a^14^, may promote glycolysis and inflammatory gene expression through HIF-1α–mediated transcriptional reprogramming.

Concomitant with the glycolytic shift, ORF7a markedly suppressed mitochondrial respiration. Mitochondrial flux analysis revealed reduced basal and maximal respiration, diminished ATP production, and loss of spare respiratory capacity (**Figure 3a-b**), consistent with impaired mitochondrial function observed in SARS-CoV-2-infected human monocytes^11^ and PBMC from COVID19 patients^10^. Importantly, these defects occurred without detectable changes in mitochondrial structure (**Figure 3c-d**) or ETC protein abundance (**Figure S3**), indicating a functional uncoupling of respiration rather than a loss of mitochondrial content. Flow cytometry confirmed a decrease in mitochondrial membrane potential (ΔΨm) (**Figure 3e-f**), further supporting a collapse in bioenergetic function. Loss of ΔΨm is a hallmark of mitochondrial dysfunction in SARS-CoV-2 infection and has been previously observed in response to other viral accessory proteins such as ORF3a, which triggers mitochondrial depolarization and apoptosis through ROS accumulation^22,24^.

ROS are at the nexus of chronic inflammation and metabolic regulation, critically driving mitochondrial oxidative phosphorylation, inflammatory cytokine activation and tissue damage^18,43^. ROS are produced metabolically by mitochondria as part of the respiratory electron transport chain, and under physiological conditions, ROS generation and antioxidant defenses are tightly balanced. However, ROS overproduction disrupts this equilibrium, triggering oxidative stress and potential cell death^31^. Increased oxidative stress in severe COVID-19 contributes to inflammation, endothelial cell dysfunction, thrombosis that can lead to multiorgan damage^44^. We detected that ORF7a induces a pronounced oxidative stress phenotype, as demonstrated by reduced levels of oxidized (GSSG) and reduced glutathione (GSH) (**Figure 1e**, **Table 1**), together with an increase in ROS levels detected by flow cytometry (**Figure 4b**) and alteration in key antioxidant enzymes (**Figure 4e-f**). These findings suggest that ORF7a disrupts redox homeostasis by elevating mitochondrial ROS production while dampening the antioxidant capacity of the cell, a combination that may sensitize cells to oxidative damage and contribute to tissue pathology during infection.

Proteomic data also highlighted upregulation of CPS1 (**Figure 1a**), a key urea cycle enzyme which catalyzes the first and rate-limiting step of the urea cycle, playing a critical role in nitrogen metabolism by converting ammonia and bicarbonate into carbamoyl phosphate using ATP^45^. This process is essential for detoxifying excess nitrogen derived from amino acid catabolism, particularly under conditions of metabolic stress or mitochondrial dysfunction. The marked upregulation of CPS1 observed in ORF7a-expressing cells may reflect a compensatory mechanism to buffer intracellular nitrogen imbalance, possibly arising from increased amino acid turnover or mitochondrial stress-induced proteolysis. In hepatocytes, CPS1 is tightly coupled to mitochondrial bioenergetics and is sensitive to alterations in mitochondrial membrane potential and redox state^46,47^.

ORF7a also enhanced the expression of enzymes involved in fatty acid β-oxidation and lipid activation, such as ACDVL, HADHA, and ACSL, possibly to support the energy demands of glycolysis-dependent cells or to enable lipid remodeling required for viral replication (**Figure 1d-e**). Supporting this, metabolomic analysis revealed accumulation of lysophosphatidylethanolamines and unsaturated fatty acids, indicative of membrane turnover and lipid signaling (**Figure 1e**, **Table 1**). ORF3a has been shown to be both necessary and sufficient to induce lipid droplets accumulation during SARS-CoV-2 infection, thereby promoting efficient virus replication^48^.

On the other hand, the observed downregulation of TALDO1 and PRPS1 in A549-ORF7a cells suggests a targeted disruption of the PPP and de novo nucleotide biosynthesis (**Figure 1e-d**). TALDO1 plays a central role in the non-oxidative branch of the pentose phosphate pathway, which provides ribose-5-phosphate for nucleotide synthesis and maintains redox balance through NADPH production^49^. PRPS1 catalyzes the formation of phosphoribosyl pyrophosphate (PRPP), a key precursor for purine and pyrimidine nucleotide biosynthesis^50^. The concomitant reduction in nucleotide monophosphate levels (UMP, AMP, GMP, dGMP) observed in our metabolomic analysis (**Figure 1e**, **Table 1**) indicates a functional consequence of this metabolic reprogramming, likely to impair DNA/RNA synthesis and cellular proliferation. This phenotype could contribute to the antiviral effects of ORF7a by restricting host cell proliferation or reflect a viral strategy to modulate nucleotide availability to favor viral RNA synthesis. Further investigation is warranted to clarify whether these alterations result from direct ORF7a–host protein interactions or arise as part of a broader cellular stress response.

A key mechanistic insight was the upregulation of PDK4, a kinase that phosphorylates and inactivates the PDH complex, in response to ORF7a (**Figure 5d-f**). This was supported by increased PDHE1α phosphorylation, a hallmark of PDH inhibition^33,38^, in both A549 (**Figure 5e**) and THP1-MΦ cells (**Figure 5g**). The upregulation of PDK4 by ORF7a and the resulting inhibition of the PDH complex point to a strategic metabolic reprogramming that limits mitochondrial pyruvate oxidation. Beyond its role in viral pathogenesis, PDK4 has been extensively studied in the context of metabolic diseases such as type 2 diabetes, where its overexpression contributes to impaired glucose oxidation, increased lipid utilization, and insulin resistance^36–38,51^. In diabetic individuals, elevated PDK4 expression exacerbates hyperglycemia by reducing PDH activity, thereby decreasing glucose utilization and promoting gluconeogenesis^51^. The induction of PDK4 by ORF7a may therefore have profound consequences in patients with pre-existing metabolic disorders, potentially worsening insulin resistance and glycemic control during SARS-CoV-2 infection. This is particularly relevant given that diabetes is a major risk factor for severe COVID-19 and is associated with poor clinical outcomes^12,52^. Moreover, pharmacological inhibitors of PDKs have been proposed as potential therapeutic agents to restore mitochondrial function in COVID-19 and other conditions involving metabolic dysfunction^53,54^. We used DCA to inhibit PDK4 and restore PDH activity (**Figure 6**). While DCA successfully decreased PDHE1α phosphorylation and increased mitochondrial respiration in control cells, it failed to rescue OXPHOS in ORF7a-expressing cells. In fact, maximal respiration further declined upon DCA treatment, indicating that mitochondrial impairment extends beyond PDH inhibition. These results suggest that ORF7a disrupts mitochondrial respiration through additional, PDH-independent mechanisms.

Intriguingly, despite the inhibition of the PDH complex by ORF7a-induced *PDK4* expression, metabolomic analysis revealed an accumulation of alanine rather than lactate (**Figure 1e**). This suggests a preferential transamination of pyruvate to alanine, potentially via alanine aminotransferase (ALT), as a compensatory mechanism to limit cytosolic pyruvate accumulation, minimize redox stress, and sustain NAD⁺ recycling. This diversion may represent a unique metabolic signature of ORF7a, differing from the classic Warburg phenotype seen in cancer and other viral infections^55–57^.

Subsequent high-resolution respirometry in permeabilized cells identified complex I as a selective target of ORF7a-mediated inhibition. ORF7a-expressing cells displayed a blunted OCR response to NADH, but normal responses to complex II (succinate) and complex IV (TMPD/ascorbate) substrates, indicating a specific impairment of complex I activity (**Figure 7**). Crucially, assays in isolated mitochondria confirmed these findings and excluded the involvement of alternative NADH-consuming enzymes, establishing complex I as a direct target of ORF7a (**Figure 7d**). BN-PAGE further demonstrated reduced incorporation of complex I into higher-order supercomplexes (**Figure 7e-f**), linking enzymatic dysfunction to defective supramolecular organization of the ETC.

Complex I is the largest and most intricate ETC complex, and its assembly is highly sensitive to mitochondrial stress, proteostasis, and membrane potential^58^. Disruption of this process, whether due to oxidative stress, mtDNA damage, or interference with assembly factors, can lead to decreased enzymatic activity despite the presence of intact subunits^59^. Virally encoded proteins, including those from SARS-CoV-2, have been shown to localize to mitochondria and alter organelle dynamics, translation, and import, all of which are critical for ETC complex formation^22,60–62^.

In particular, the observed mitochondrial stress signature in ORF7a-expressing cells, characterized by PDH inhibition, ROS imbalance, and failure to restore respiration after DCA treatment, may impair the assembly of complex I by affecting mitochondrial proteostasis or membrane potential, both of which are required for efficient insertion and stabilization of complex I subunits.

This selective complex I disruption is functionally significant, as complex I is the primary entry point of electrons into the ETC and a major source of mitochondrial ROS. The observed increase in intracellular ROS, coupled with depletion of glutathione and dipeptide antioxidants, suggests that ORF7a-induced complex I dysfunction leads to oxidative stress, further impairing mitochondrial function and redox homeostasis.

Taken together, our findings demonstrated that ORF7a acts at multiple regulatory nodes to reprogram host metabolism, including enhancing glycolysis, inhibiting PDH, destabilizing complex I and impairing antioxidant defenses. This convergence drives a glycolytic, redox-stressed phenotype that may promote viral replication and persistence while contributing to host tissue damage. Importantly, by linking enzymatic inhibition with defective supercomplex assembly, our study identifies a structural and functional vulnerability of the mitochondrial respiratory chain to viral manipulation. As APs such as ORF7a are often overlooked in therapeutic development, targeting host mitochondrial pathways, particularly complex I and its supramolecular organization, may represent a promising strategy to mitigate the metabolic dysregulation associated with SARS-CoV-2 infection.

## Material and Methods

### Cell culture, lentivirus production and transduction

A549 pulmonary epithelial cells (ATCC CRM-CCL-185; RRID: CVCL_0023) were cultured in Dulbecco’s Modified Eagle Medium (DMEM) (Gibco, #41966029) supplemented with 10% (v/v) heat-inactivated fetal bovine serum (FBS) (Gibco, #1027016) and 1% Penicillin-Streptomycin (100U/ml) (Gibco, #15070063). THP1 (ATCC CRM-TIB-202; RRID: CVCL_0006) were cultured in Roswell Park Memorial Institute (RPMI) (Gibco, #11875093) supplemented with 10% (v/v) heat-inactivated fetal bovine serum (FBS) (Gibco, #1027016) and 1% Penicillin-Streptomycin (100U/ml) (Gibco, #15070063). ORF7a accessory protein coding sequences were cloned into pLVX-EF1α-IRES-Puro Cloning and Expression Lentivector (Clontech, Takara, #631253) to generate pseudotyped lentiviral particles encoding ORF7a protein (Wuhan-Hu-1 isolate) as described previously ^30^. Cells were transduced by incubating them with lentivirus at a MOI of 10 for 24 h followed by 2 µg/ml puromycin treatment to start the selection of successfully transduced cells. Differentiation of THP1 into macrophage-like cells (THP1-MƟ) was induced by treating 80 ng/ml PMA phorbol 12-myristate 13-acetate (PMA) (Sigma-Aldrich, P8139) for 24 hours. After PMA exposure, the cells were washed with PBS and incubated with fresh medium for an additional 24–72 hours prior to experiments to allow complete differentiation. All cells were cultured at 37°C in a 5% CO2, 90% humidity atmosphere.

### Protein Sample Preparation and Proteomics

Cells were harvested and lysed in ice-cold RIPA buffer. Protein concentration was measured using the Pierce™ BCA Protein Assay Kits (Thermo Scientific Inc., Waltham, MA, USA), and 100 µg were sent to the Proteomics Facility at Research Support Central Service, University of Cordoba (SCAI-UCO), for sample preparation and analysis. Protein extracts underwent clean-up in one dimensional sodium dodecyl sulfate-polyacrylamide gel electrophoresis with 10% polyacrylamide. After electrophoresis at 100 V, gel was stained with Coomassie Blue and protein bands were cut out and stored in water until digestion. Protein bands were first destained in 200 mM ammonium bicarbonate (AB)/50% acetonitrile, then in 100% acetonitrile, reduced with 20 mM dithiothreitol and incubated in 25 mM AB at 55 °C. After cooling to room temperature, free thiols were alkylated with 40 mM iodoacetamide. Following two AB washes, proteins were digested with trypsin (Promega, Madison, WI, USA) and incubated overnight at 37 °C. Digestion was stopped by adding trifluoroacetic acid to a final concentration of 1%. Digested samples were dried and resuspended in 0.1% formic acid (FA).

Nano liquid chromatography (LC) was performed on a Dionex Ultimate 3000 nano ultra performance LC (Thermo Scientific Inc., Waltham, MA, USA) with a C18 75 µm × 50 Acclaim Pepmap column (Thermo Scientific Inc., Waltham, MA, USA). Peptides were loaded onto a 300 µm × 5 mm Acclaim Pepmap precolumn (Thermo Scientific Inc., Waltham, MA, USA) in 2% acetonitrile/0.05% trifluoroacetic acid, then separated at 40 °C with a gradient of buffer A (water with 0.1% FA) and buffer B (20% acetonitrile with 0.1% FA) over 85 min of chromatography. Peptide cations were ionized and analyzed on a Thermo Orbitrap Fusion (Q-OT-qIT, Thermo Scientific Inc., Waltham, MA, USA), operated in positive mode with survey scans at 120K resolution. Tandem mass spectrometry (MS/MS) involved quadrupole isolation, collision-induced dissociation fragmentation, and rapid scan MS analysis, sampling only precursors with charge states 2–5. The instrument operated in top speed mode with 3 s MS2 cycles, excluding precursors dynamically.

MS2 spectra were analyzed using MaxQuant software v. 1.6.17.0^63^, with the Andromeda search engine set against a Uniprot_proteome_Homo-sapiens_database. Peptides were generated through tryptic digestion with up to one missed cleavage, fixed carbamidomethylation of cysteines, and variable oxidation of methionine. The mass tolerance was 10 ppm, and product ions were searched with 0.6 Da tolerance. Peptide matches were filtered to a 1% false discovery rate (FDR). Quantification was conducted using the MaxLFQ label-free method^64^, using retention time alignment and match-between-runs protocol.

### Untargeted metabolomic

Metabolites were extracted by methanol precipitation and analyzed by liquid chromatography–mass spectrometry (LC–MS) using a Bruker Elute UHPLC system coupled to a TIMS-TOF Pro instrument operating in both positive and negative ion modes with 4D PASEF acquisition. Chromatographic separation was achieved on a C18 reversed-phase column with a 30-min gradient of water and acetonitrile, both containing 0.1% formic acid. Calibration and online recalibration were performed using sodium formate clusters to ensure mass accuracy. Raw data were processed with DataAnalysis 6.1 (Bruker Daltonics) and untargeted feature extraction was carried out in MetaboScape 8.0.2 (Bruker Daltonics) using the T-ReX algorithm for peak detection, alignment, and normalization. Quality control (QC) samples were included every 10 injections, and features were filtered based on reproducibility in pooled QC samples and response linearity in a QC dilution series (R² > 0.65). Signals from both ionization modes were merged and only features consistently detected across replicates were retained for statistical analysis.

### Real-time qPCR

RNA samples (500 ng) were reverse transcribed using qScript™ cDNA synthesis kit (Quanta Biosciences Inc., #95047), following manufacturer’s instructions. The final 15 µL PCR reaction included 2 μL of 1:5 diluted cDNA as template, 3 µL of 5x PyroTaq EvaGreen qPCR Mix Plus with ROX (Cultek Molecular Bioline, #88H24), and transcript-specific forward and reverse primers (**Table S3**) at a 20 μM final concentration. Real time PCR was carried out in a QuantStudio 12K Flex system (Applied Biosystems) under the following conditions: 15 min at 95°C followed by 40 cycles of 30 s at 94°C, 30 s at 57°C and 45 s at 72°C. Melting curve analyses were performed at the end, in order to ensure specificity of each PCR product. Relative expression results were calculated using GenEx6 Pro software (MultiD-Göteborg, Sweden), based on the Cq values obtained.

### Western blots

Transduced cells were harvested and lysed in ice-cold RIPA Lysis Buffer containing complete protease inhibitor and phosphatase inhibitors cocktails (Sigma-Aldrich) at 4° C. Cell lysates were mixed with 5× SDS-PAGE Sample Loading Buffer, and heated at 95° C for 5 min. Protein samples were resolved by SDS polyacrylamide gel electrophoresis and transferred onto a PVDF membrane using Trans-Blot Turbo Transfer System (Bio-Rad, Hercules, CA), followed by blocking for 1 h with 5% nonfat milk in Tris-buffered saline-Tween 20 buffer (TBST) and probing with corresponding primary and secondary antibodies (**Table S4**). The proteins were visualized by chemoluminiscence using ChemiDoc Imaging Systems (Bio-Rad). Relative protein expression was calculated by sequentially normalizing against the loading control (GAPDH).

### Transmission electron microscopy

Cell monolayers were washed with PBS 1× and fixed in situ for 1 h at room temperature (RT) with 3% glutaraldehyde (EM Grade, Ted Pella INC) in PBS 1×. Fixed cells were washed three times with PBS 1×. Cell postfixation was as follows: 1 h at 4× with 1% osmium tetroxide (Electron Microscopy Sciences) and 0,8% potassium ferricyanide in PBS, washed with PBS three times, and after dehydration in a gradient of 30% to 100% in ETOH, cells were embedded in a gradient of EtOH/LX 112 epoxy resin to 100% epoxy resin (Ladd Research). The samples were polymerized at 60°C for 2 days. Ultrathin 70 nm-thick sections were obtained with a Leica EM UC6 ultramicrotome (Leica Microsystems GmbH), transferred to collodion/carbon-coated EM grids and stained with Uranyl Acetate 5% for 20 min and Reynold’s Lead Citrate for 5 min. 70 nm sections were visualized on a FEI Tecnai 12 electron microscope equipped with a LaB6 filament and operated at 100 kV. Images were recorded with a FEI Ceta digital camera at various magnifications. Median of mitochondria longest diameter was measured with ImageJ program, and median of ridges was counted in high resolution ME images. In both cases, at least 50 mitochondria were analyzed.

### Mitochondrial membrane potential (ΔΨm)

Cells were seeded at 300,000 cells/well in 6-well plates 24 h before staining. For quantitative analysis of mitochondrial membrane potential (ΔΨm), A549 cells were stained in DMEM 1% penicillin/streptomycin with 5 nM MitoView 633 (Biotium, #70055). Plates were incubated at 37℃ for 30 min protected from light. Cells were then harvested, washed with PBS 1× and resuspended in PBS. For these experiments, 10,000 events were analysed using a CytoFLEX flow cytometer (Beckman Coulter) and FlowJo v10 software (BD Biosciences).

### Reactive oxygen species (ROS) analysis

ROS production was evaluated using the ROS Detection Assay Kit (Canvax Biotec S.L, Cordoba, Spain) which contains the cell-permeant reagent dichlorodihydrofluorescein diacetate (H_2_DCFDA), an indicator of reactive oxygen intermediates, that becomes fluorescent when oxidized, following the manufacturer’s instructions. Cells were seeded on a 24 well plates at 7,5x10^4^ cells per well 2-day prior assay. Cells were incubated with H_2_DCFDA (25 mM) for 45 minutes at 37°C. Positive control cells were treated with 100 µM H_2_O_2_. ROS were measured by flow cytometry (BD Accuri C6 Plus Flow Cytometer).

### Seahorse extracellular flux analysis

Agilent Seahorse XF assays were performed to determine oxygen consumption rate (OCR) and extracellular acidification rate (ECAR) using an Agilent Seahorse XF24 Analyzer (Seahorse Bioscience, Agilent), previously calibrated overnight in a CO₂-free incubator with Seahorse XF Calibrant solution. 24 h before the assay, a total of 37,500 A549 cells and 200,000 THP1-MƟ cells were seeded in a Seahorse 24-well XF Cell Culture Microplate in culture medium. The cells were then allowed to adhere for 24 hours in a 5% CO₂ atmosphere at 37 °C.

On the day of assay, for Agilent Seahorse MitoStress Test media was changed to XF DMEM medium supplemented with 10 mM glucose, 1 mM sodium pyruvate and 2 mM glutamine, pH 7.4, and then maintained in XF assay media at 37 °C in a CO_2_-free incubator 1 h. Mitochondrial function of the cells was analyzed by sequential injections of the modulators oligomycin (1 μM), carbonyl cyanide-*p*-trifluoromethoxyphenylhydrazone (FCCP, 1 μM) and a mixture of antimycin A and rotenone (Rot/AA, 0.5 μM). This assay allows measurement of key parameters of mitochondrial respiration including basal respiration, ATP production, maximal respiration, spare respiratory capacity, proton leak, and non-mitochondrial respiration.

For the Agilent Seahorse Glycolysis Stress Test, the cell culture medium was replaced with XF DMEM medium supplemented with 2 mM glutamine, pH 7.4, and cells were incubated in XF assay media at 37 °C in a CO₂-free incubator for 1 hour. The glycolytic function of the cells was analyzed by sequential injections of glucose (10 mM), oligomycin (1 μM), and 2-deoxy-D-glucose (2-DG, 50 mM), allowing measurement of glycolysis, glycolytic capacity, and glycolytic reserve, respectively. Data were collected using Agilent Seahorse Wave 2.6.1 Desktop software and normalized to protein concentration determined at the end of the assay. Data were exported to GraphPad Prism version 9 for analysis.

### Respirometry In Frozen Samples (RIFS) Assay

The oxygen consumption rate of THP1-derived macrophage (THP1-MƟ) cells was measured using a recently developed method permitting respirometry analysis of frozen cells and mitochondrial fractions isolated from previously frozen cells^65^. THP1 cells were seeded into Seahorse XF24 plates at a density of 200.000 cells per well and subjected to three freeze-thaw cycles with intermediate washes in MAS buffer (70 mM sucrose, 220 mM mannitol, 5 mM KH2PO4, 5 mM MgCl2, 1 mM EGTA, 2 mM HEPES, pH 7,4) supplemented with protease inhibitors. Following these washes, cells were resuspended in 50 μl of MAS buffer supplemented with protease inhibitors and cytochrome c. The plates were then centrifuged at 2.300 g for 5 min at 4°C (no brake), and 450 μl of MAS was carefully added to each well prior to respirometry measurements (alamethicin was added to study NADH oxidation).

The mitochondrial fraction was isolated from THP1 cells as an alternative approach to assess cellular respiration. Briefly, THP1 cells were centrifuged at 700 g for 5 min at room temperature, and the resulting pellet was frozen until further processing. Cells were mechanically homogenized with Tyssuelyser (Qiagen), the homogenates were centrifuged at 1.000 g for 5 min at 4°C; then, the supernatant was collected and centrifuged at 12.000 g for 12 min at 4°C. The pellet was resuspended in Extraction Medium. Protein concentration was determined by BCA (Thermo Fisher). Mitochondrial fractions (15 μg) were loaded into Seahorse XF96 microplate in 20 μl of MAS containing cytochrome c (10 μg/ml, final concentration). The loaded plate was centrifuged at 2,000 g for 5 min at 4°C (no brake), and an additional 130 μl of MAS + cytochrome c was added to each well.

Mitochondrial respiration linked to NADH oxidation was stimulated by the addition of NADH to promote electron flow through complexes I, III, and IV. Specificity of this respiratory activity was confirmed by subsequent inhibition with antimycin A. Following this, Complex IV activity was assessed by stimulating oxygen consumption with N,N,N′,N′-tetramethyl-p-phenylenediamine (TMPD) in combination with ascorbic acid, and subsequently inhibiting respiration with sodium azide.

In a separate experimental condition, succinate was added in the presence of rotenone to inhibit NADH-linked pathways and stimulate electron flow through the succinate oxidation pathway involving complexes II, III, and IV. Oxygen consumption under these conditions was measured and then inhibited by antimycin A. Complex IV activity was evaluated as described above using TMPD and ascorbic acid to stimulate respiration, followed by sodium azide inhibition.

### Blue Native PAGE (BN-PAGE)

Mitochondrial supercomplex organization was analyzed by Blue Native gel electrophoresis following established protocols^66^. Isolated mitochondrial fractions (75 µg) were solubilized with digitonin to preserve respiratory supercomplexes and subsequently separated by BN-PAGE. Individual complexes and their supramolecular assemblies (respirasomes) were visualized and quantified by immunoblotting with antibodies against representative subunits of complexes I, III, and IV (**Table S4**).

## Acknowledgments

This research work was funded by the European Commission – NextGenerationEU (Regulation EU 2020/2094) through CSIC’s Global Health Platform (PTI+ Salud Global) (COVID-19-117 and SGL2103015), Junta de Andalucía (CV20-20089) and Spanish Ministry of Science projects (PID2021-123399OB-I00 and PID2024-162356OB-I00). TGG is recipient of a Ramón y Cajal contract (RYC2021-031614-I) funded by MCIN/AEU/10.13039/501100011033 and NextGeneration EU/PRTR. RFR is recipient of a Contrato Predoctoral para Personal Investigador en Formación (PIF), University of Córdoba. We would like to thank Eduardo Chicano-Galvez and Ángela Peralbo-Molina from IMIBIC Mass Spectrometry and Molecular Imaging Unit (IMSMI) for performing the metabolomic analysis and Carlos Fuentes-Almagro from the Proteomics Facility at Research Support Central Service, University of Cordoba (SCAI-UCO), for sample preparation and proteomic analysis.

## Author contributions

Conceptualization: T.G.G and J.J.G. Methodology: R.F.R, C.M.S.J, R.A.P, L.M.S.M, A.D.P, B.D.L.A and T.G.G. Investigation: All authors. Writing original draft: T.G.G. Writing-review and editing: J.J.G, M.M, J.M.V, J.A.E. and T.G.G. Funding acquisition: J.J.G, M.M and T.G.G.

## References

1. Batabyal, R. et al. Metabolic dysfunction and immunometabolism in COVID-19 pathophysiology and therapeutics. Int. J. Obes. 45, 1163–1169 (2021).

2. Steenblock, C. et al. COVID-19 and metabolic disease: mechanisms and clinical management. Lancet Diabetes Endocrinol. 9, 786–798 (2021).

3. Wu, D. et al. Plasma metabolomic and lipidomic alterations associated with COVID-19. Natl. Sci. Rev. 7, 1157–1168 (2020).

4. Bruzzone, C. et al. SARS-CoV-2 Infection Dysregulates the Metabolomic and Lipidomic Profiles of Serum. iScience 23, 101645 (2020).

5. Sanchez, E. L. & Lagunoff, M. Viral activation of cellular metabolism. Virology 479–480, 609–618 (2015).

6. Bappy, S. S. et al. Virus-induced host cell metabolic alteration. Rev. Med. Virol. 34, e2505 (2024).

7. Palmer, C. S. Innate metabolic responses against viral infections. Nat. Metab. 4, 1245–1259 (2022).

8. Allen, C. N. S. et al. SARS-CoV-2 Causes Lung Inflammation through Metabolic Reprogramming and RAGE. Viruses 14, 983 (2022).

9. Andrade Silva, M., da Silva, A. R. P. A., do Amaral, M. A., Fragas, M. G. & Câmara, N. O. S. Metabolic Alterations in SARS-CoV-2 Infection and Its Implication in Kidney Dysfunction. Front. Physiol. 12, 624698 (2021).

10. Ajaz, S. et al. Mitochondrial metabolic manipulation by SARS-CoV-2 in peripheral blood mononuclear cells of patients with COVID-19. Am. J. Physiol.-Cell Physiol. 320, C57–C65 (2021).

11. Codo, A. C. et al. Elevated Glucose Levels Favor SARS-CoV-2 Infection and Monocyte Response through a HIF-1α/Glycolysis-Dependent Axis. Cell Metab. 32, 437–446.e5 (2020).

12. Bornstein, S. R. et al. Practical recommendations for the management of diabetes in patients with COVID-19. Lancet Diabetes Endocrinol. 8, 546–550 (2020).

13. Meng, X. et al. HIF-1α promotes virus replication and cytokine storm in H1N1 virus-induced severe pneumonia through cellular metabolic reprogramming. Virol. Sin. 39, 81–96 (2024).

14. Tian, M. et al. HIF-1α promotes SARS-CoV-2 infection and aggravates inflammatory responses to COVID-19. Signal Transduct. Target. Ther. 6, 308 (2021).

15. Zhang, Y. et al. Influenza A virus-induced glycolysis facilitates virus replication by activating ROS/HIF-1α pathway. Free Radic. Biol. Med. 225, 910–924 (2024).

16. Ren, Z. et al. The Triangle Relationship Between Long Noncoding RNA, RIG-I-like Receptor Signaling Pathway, and Glycolysis. Front. Microbiol. 12, 807737 (2021).

17. Chan, D. C. Mitochondria: dynamic organelles in disease, aging, and development. Cell 125, 1241–1252 (2006).

18. Burtscher, J., Cappellano, G., Omori, A., Koshiba, T. & Millet, G. P. Mitochondria: In the Cross Fire of SARS-CoV-2 and Immunity. iScience 23, 101631 (2020).

19. Shin, H. J. et al. SARS-CoV-2 aberrantly elevates mitochondrial bioenergetics to induce robust virus propagation. Signal Transduct. Target. Ther. 9, 125 (2024).

20. Guarnieri, J. W., et al. Mitochondrial antioxidants abate SARS-COV-2 pathology in mice. Proc. Natl. Acad. Sci. 121, e2321972121 (2024).

21. Jana, S. et al. HIF-1α-Dependent Metabolic Reprogramming, Oxidative Stress, and Bioenergetic Dysfunction in SARS-CoV-2-Infected Hamsters. Int. J. Mol. Sci. 24, 558 (2022).

22. Qudus, M. S. et al. SARS-CoV-2-ORF-3a Mediates Apoptosis Through Mitochondrial Dysfunction Modulated by the K+ Ion Channel. Int. J. Mol. Sci. 26, 1575 (2025).

23. Redondo, N., Zaldívar-López, S., Garrido, J. J. & Montoya, M. SARS-CoV-2 Accessory Proteins in Viral Pathogenesis: Knowns and Unknowns. Front. Immunol. 12, 708264 (2021).

24. López-Ayllón, B. D. et al. Metabolic and mitochondria alterations induced by SARS-CoV-2 accessory proteins ORF3a, ORF9b, ORF9c and ORF10. J. Med. Virol. 96, e29752 (2024).

25. Zhang, J. et al. Understanding the Role of SARS-CoV-2 ORF3a in Viral Pathogenesis and COVID-19. Front. Microbiol. 13, 854567 (2022).

26. Arshad, N. et al. SARS-CoV-2 accessory proteins ORF7a and ORF3a use distinct mechanisms to down-regulate MHC-I surface expression. Proc. Natl. Acad. Sci. U. S. A. 120, e2208525120 (2023).

27. Liu, Z. et al. Ubiquitination of SARS-CoV-2 ORF7a Prevents Cell Death Induced by Recruiting BclXL To Activate ER Stress. Microbiol. Spectr. 10, e0150922 (2022).

28. Cao, Z. et al. Ubiquitination of SARS-CoV-2 ORF7a promotes antagonism of interferon response. Cell. Mol. Immunol. 18, 746–748 (2021).

29. Timilsina, U., Umthong, S., Ivey, E. B., Waxman, B. & Stavrou, S. SARS-CoV-2 ORF7a potently inhibits the antiviral effect of the host factor SERINC5. Nat. Commun. 13, 2935 (2022).

30. García-García, T. et al. Impairment of antiviral immune response and disruption of cellular functions by SARS-CoV-2 ORF7a and ORF7b. iScience 25, 105444 (2022).

31. Zorov, D. B., Juhaszova, M. & Sollott, S. J. Mitochondrial Reactive Oxygen Species (ROS) and ROS-Induced ROS Release. Physiol. Rev. 94, 909–950 (2014).

32. Jeong, J. Y., Jeoung, N. H., Park, K.-G. & Lee, I.-K. Transcriptional regulation of pyruvate dehydrogenase kinase. Diabetes Metab. J. 36, 328–335 (2012).

33. Sugden, M. C. & Holness, M. J. Mechanisms underlying regulation of the expression and activities of the mammalian pyruvate dehydrogenase kinases. Arch. Physiol. Biochem. 112, 139–149 (2006).

34. Pettersen, I. K. N. et al. Upregulated PDK4 expression is a sensitive marker of increased fatty acid oxidation. Mitochondrion 49, 97–110 (2019).

35. Liu, B., Zhang, Y. & Suo, J. Increased Expression of PDK4 Was Displayed in Gastric Cancer and Exhibited an Association With Glucose Metabolism. Front. Genet. 12, (2021).

36. Lee, I.-K. The Role of Pyruvate Dehydrogenase Kinase in Diabetes and Obesity. Diabetes Metab. J. 38, 181–186 (2014).

37. Kim, Y. I., Lee, F. N., Choi, W. S., Lee, S. & Youn, J. H. Insulin regulation of skeletal muscle PDK4 mRNA expression is impaired in acute insulin-resistant states. Diabetes 55, 2311–2317 (2006).

38. Park, S. et al. Role of the Pyruvate Dehydrogenase Complex in Metabolic Remodeling: Differential Pyruvate Dehydrogenase Complex Functions in Metabolism. Diabetes Metab. J. 42, 270–281 (2018).

39. Schoenmann, N., Tannenbaum, N., Hodgeman, R. M. & Raju, R. P. Regulating mitochondrial metabolism by targeting pyruvate dehydrogenase with dichloroacetate, a metabolic messenger. Biochim. Biophys. Acta BBA - Mol. Basis Dis. 1869, 166769 (2023).

40. Thompson, E. A. et al. Metabolic programs define dysfunctional immune responses in severe COVID-19 patients. Cell Rep. 34, 108863 (2021).

41. Moolamalla, S. T. R., Balasubramanian, R., Chauhan, R., Priyakumar, U. D. & Vinod, P. K. Host metabolic reprogramming in response to SARS-CoV-2 infection: A systems biology approach. Microb. Pathog. 158, 105114 (2021).

42. Mullen, P. J. et al. SARS-CoV-2 infection rewires host cell metabolism and is potentially susceptible to mTORC1 inhibition. Nat. Commun. 12, 1876 (2021).

43. Mittal, M., Siddiqui, M. R., Tran, K., Reddy, S. P. & Malik, A. B. Reactive Oxygen Species in Inflammation and Tissue Injury. Antioxid. Redox Signal. 20, 1126–1167 (2014).

44. Alam, M. S. & Czajkowsky, D. M. SARS-CoV-2 infection and oxidative stress: Pathophysiological insight into thrombosis and therapeutic opportunities. Cytokine Growth Factor Rev. 63, 44–57 (2022).

45. Nitzahn, M. & Lipshutz, G. S. CPS1: Looking at an Ancient Enzyme in a Modern Light. Mol. Genet. Metab. 131, 289–298 (2020).

46. Li, P., Kuo, N., Patel, R. & Omary, M. B. Hypoosmosis alters hepatocyte mitochondrial morphology and induces selective release of carbamoyl phosphate synthetase 1. Am. J. Physiol. Gastrointest. Liver Physiol. 325, G334–G346 (2023).

47. Nakagawa, T. & Guarente, L. Urea cycle regulation by mitochondrial sirtuin, SIRT5. Aging 1, 578–581 (2009).

48. Wang, W. et al. Genetic variety of ORF3a shapes SARS-CoV-2 fitness through modulation of lipid droplet. J. Med. Virol. 95, e28630 (2023).

49. Stincone, A. et al. The return of metabolism: biochemistry and physiology of the pentose phosphate pathway. Biol. Rev. 90, 927–963 (2015).

50. Brouwer, A. P. M. de et al. PRPS1 Mutations: Four Distinct Syndromes and Potential Treatment. Am. J. Hum. Genet. 86, 506–518 (2010).

51. Jeoung, N. H. Pyruvate Dehydrogenase Kinases: Therapeutic Targets for Diabetes and Cancers. Diabetes Metab. J. 39, 188–197 (2015).

52. Zhou, F., et al. Clinical course and risk factors for mortality of adult inpatients with COVID-19 in Wuhan, China: a retrospective cohort study. The Lancet 395, 1054–1062 (2020).

53. Stakišaitis, D. et al. Effects of Combined Treatment with Sodium Dichloroacetate and Sodium Valproate on the Genes in Inflammation- and Immune-Related Pathways in T Lymphocytes from Patients with SARS-CoV-2 Infection with Pneumonia: Sex-Related Differences. Pharmaceutics 16, 409 (2024).

54. James, M. O. et al. Therapeutic applications of dichloroacetate and the role of glutathione transferase zeta-1. Pharmacol. Ther. 170, 166–180 (2017).

55. Vander Heiden, M. G., Cantley, L. C. & Thompson, C. B. Understanding the Warburg Effect: The Metabolic Requirements of Cell Proliferation. Science 324, 1029–1033 (2009).

56. Li, J., Wang, Y., Deng, H., Li, S. & Qiu, H.-J. Cellular metabolism hijacked by viruses for immunoevasion: potential antiviral targets. Front. Immunol. 14, 1228811 (2023).

57. Thaker, S. K., Ch’ng, J. & Christofk, H. R. Viral hijacking of cellular metabolism. BMC Biol. 17, 59 (2019).

58. Fiedorczuk, K. & Sazanov, L. A. Mammalian Mitochondrial Complex I Structure and Disease-Causing Mutations. Trends Cell Biol. 28, 835–867 (2018).

59. Signes, A. & Fernandez-Vizarra, E. Assembly of mammalian oxidative phosphorylation complexes I–V and supercomplexes. Essays Biochem. 62, 255–270 (2018).

60. Singh, K. K., Chaubey, G., Chen, J. Y. & Suravajhala, P. Decoding SARS-CoV-2 hijacking of host mitochondria in COVID-19 pathogenesis. Am. J. Physiol.-Cell Physiol. 319, C258–C267 (2020).

61. Gordon, D. E. et al. A SARS-CoV-2 protein interaction map reveals targets for drug repurposing. Nature 583, 459–468 (2020).

62. Jiang, H.-W. et al. SARS-CoV-2 Orf9b suppresses type I interferon responses by targeting TOM70. Cell. Mol. Immunol. 17, 998–1000 (2020).

63. Cox, J. & Mann, M. MaxQuant enables high peptide identification rates, individualized p.p.b.-range mass accuracies and proteome-wide protein quantification. Nat. Biotechnol. 26, 1367–1372 (2008).

64. Cox, J. et al. Accurate proteome-wide label-free quantification by delayed normalization and maximal peptide ratio extraction, termed MaxLFQ. Mol. Cell. Proteomics MCP 13, 2513–2526 (2014).

65. Acin-Perez, R. et al. A novel approach to measure mitochondrial respiration in frozen biological samples. EMBO J. 39, e104073 (2020).

66. Acín-Pérez, R., Hernansanz-Agustín, P. & Enríquez, J. A. Chapter 7 - Analyzing electron transport chain supercomplexes. in Methods in Cell Biology (eds. Pon, L. A. & Schon, E. A.) vol. 155 181–197 (Academic Press, 2020).

