## Supplementary Figures for "SARS-CoV-2 ORF7a Drives Mitochondrial Dysfunction via PDK4 Activation and Complex I Inhibition"

### Supplementary Figure 1

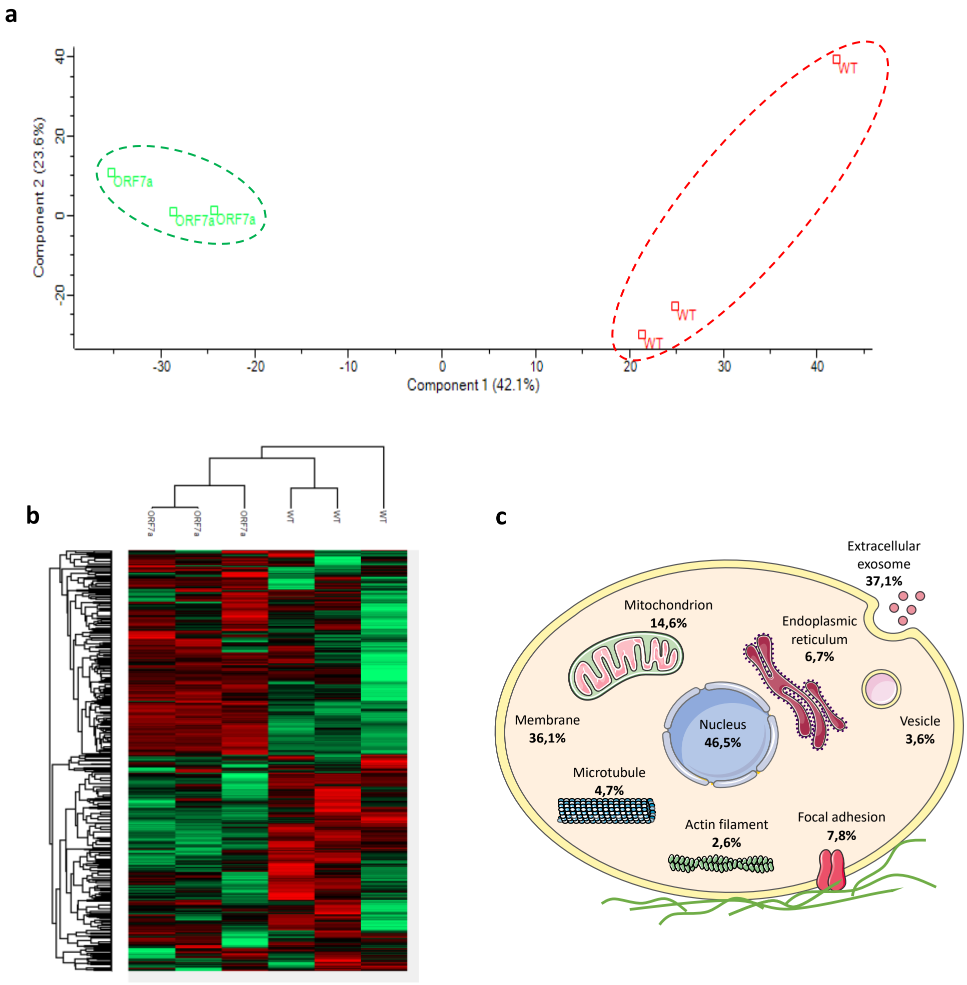


**Figure S1. Proteomics profiling of A549 cells expressing ORF7a and control cells. a.** PCA analysis of the proteome profile. Three biological replicates were generated for each sample. **b.** Heatmap analysis of all expressed proteins comparing A549-ORF7a and control cells presented with normalized protein intensities. **c.** Schematic of changes in cell components.

### Supplementary Figure 2

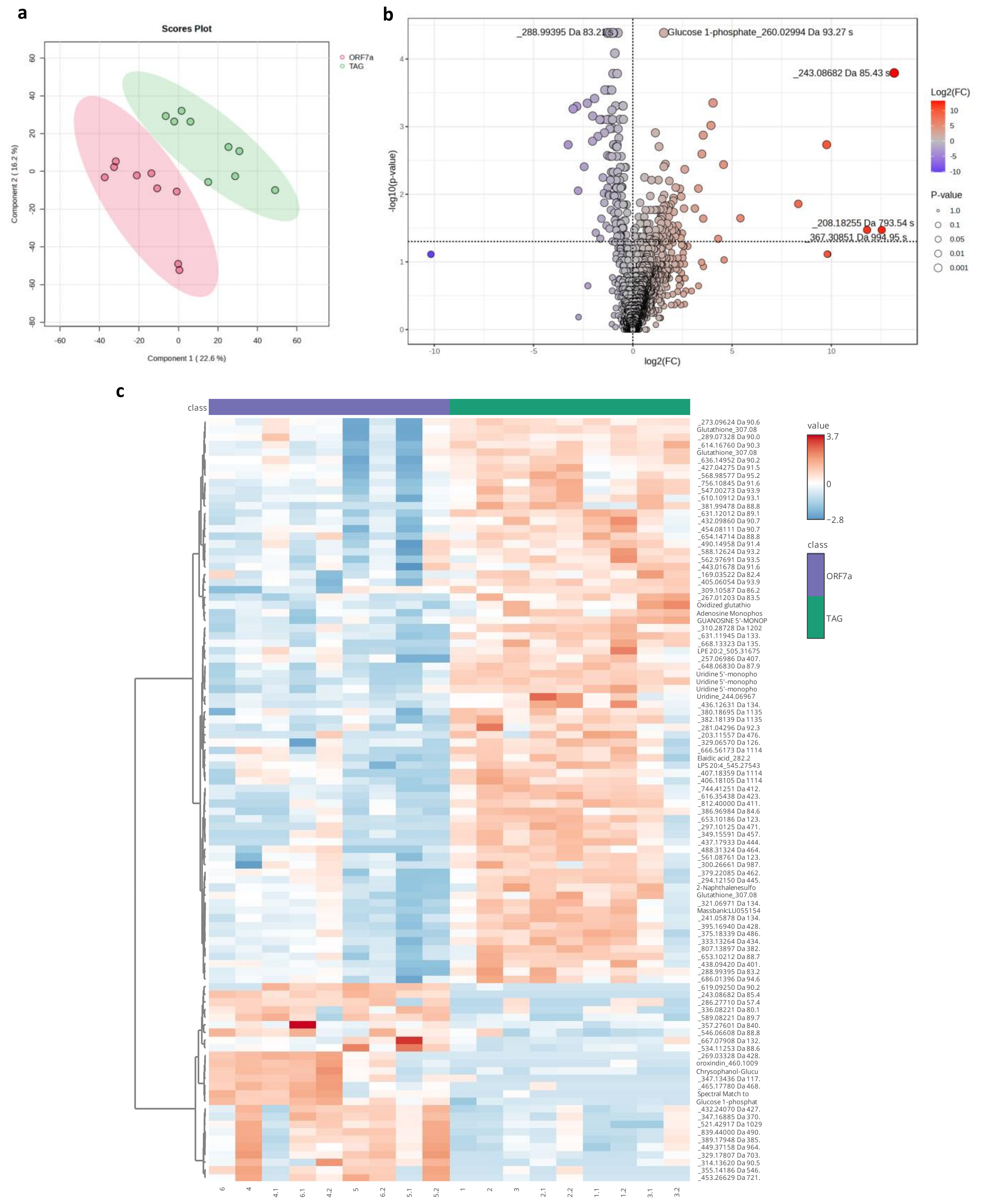


**Figure S2. Untargeted metabolomic profiling from A549 cells expressing ORF7a and control cells. a**. Partial least squares discriminant analysis (PLS-DA) of untargeted metabolomics among the two groups. **b**. volcano plots highlighted the metabolites that were increased (red) or decreased (blue) in the A549 cells expressing ORF7a. **c**. Heatmap of top 100 metabolites with p-value <0.05.

#### Supplementary Figure 3

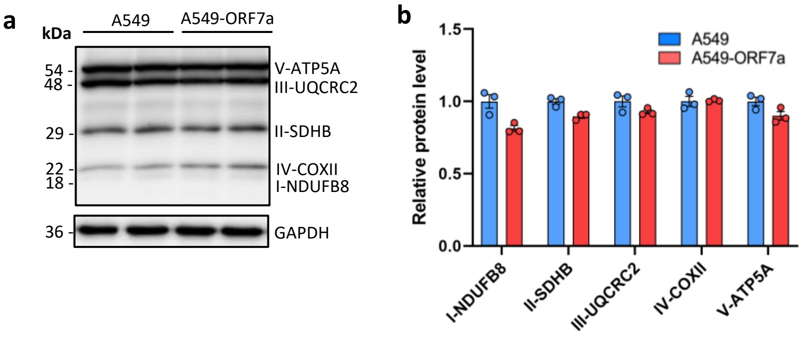


**Figure S3. Analysis of Mitochondrial OXPHOS Complex Expression by Western Blot. a**. Western blot of the indicated OXPHOS subunits obtained using human OXPHOS antibody cocktail. **b**. Densitometric quantification of band intensities corresponding to each subunit.
