## Supplementary material for "SARS-CoV-2 ORF7a Drives Mitochondrial Dysfunction via PDK4 Activation and Complex I Inhibition": Table S3

**Table S3. List of primers used for qPCR.**

| Target gen | Forward primer (5'-3') | Reverse primer (5'-3') | Reference |
|---|---|---|---|
| GLUT-1 | CTGCTCATCAACCGCAAC | CTTCTTCTCCCGCATCATCT | This work |
| HK1 | GTCTGGACGCGGGAATCTTG | CCACCACGTCCAGGTCAAAT | This work |
| PFKFB3 | CAGTTGTGGCCTCCAATATC | GGCTTCATAGCAACTGATCC | This work |
| PKM2 | ATTATTTGAGGAACTCCGCCGCCT | ATTCCGGGTCACAGCAATGATGG | This work |
| GPX1 | GTGCTCGGCTTCCCGTGCAAC | CTCGAAGAGCATGAAGTTGGGC | This work |
| PDK1 | CATGTCACGCTGGGTAATGAGG | CTCAACACGAGGTCTTGGTGCA | This work |
| PDK2 | TGCCTACGACATGGCTAAGCTC | GACGTAGACCATGTGAATCGGC | This work |
| PDK3 | TGGAAGGAGTGGGTACTGATGC | GGATTGCTCCAATCATCGGCTTC | This work |
| PDK4 | AGGTGGAGCATTTCTCGCGCTA | GAATGTTGGCGAGTCTCACAGG | This work |
| GAPDH | TGGGTGTGAACCATGAGAAG | TGGCAGTGATGGCATGGAC | This work |
