## Supplementary material for "SARS-CoV-2 ORF7a Drives Mitochondrial Dysfunction via PDK4 Activation and Complex I Inhibition": Table S4

**Table S4. List of antibodies**

| Target protein | Supplier | Catalog number |
|---|---|---|
| HIF-1a | BD | 610959 |
| PDH-E1α (pSer$^{300}$) | Sigma | AP1064 |
| PDH-E1α | Santa Cruz | Sc-377092 |
| OXPHOS cocktail | Invitrogen | 48-8199 |
| Complex I subunit NDUFA9 | Abcam | ab14713 |
| Complex II 70 kDa Fp subunit / SDHA | Invitrogen | 459200 |
| UQCRC2CORE2 | Proteintech | 14742-1-AP |
| GAPDH | Sigma | Cat#G8795; RRID:AB_1078991 |
| Anti-mouse IgG StarBrigth Blue 700 | BIO RAD | Cat.#12004158 |
| Anti-rabbit IgG StarBrigth Blue 700 | BIO RAD | Cat.#12004162 |
| Anti-mouse-HRP | Sigma | Cat#A9044; RRID:AB_258431 |
